# Small Molecule Properties Define Partitioning into Biomolecular Condensates

**DOI:** 10.1101/2022.12.19.521099

**Authors:** Sabareesan Ambadi Thody, Hanna D. Clements, Hamid Baniasadi, Andrew S. Lyon, Matthew S. Sigman, Michael K. Rosen

**Affiliations:** Department of Biophysics, Howard Hughes Medical Institute, UT Southwestern Medical Center, Dallas, TX, 75390, USA; Department of Chemistry, University of Utah, 315 South 1400 East, Salt Lake City, Utah 84112, USA; Department of Biochemistry, UT Southwestern Medical Center, Dallas, TX, 75390, USA

## Abstract

Biomolecular condensates regulate cellular function by compartmentalizing molecules without a surrounding membrane^1,2^. Condensate functions are believed to arise from the specific exclusion or enrichment of molecules^3-5^. Thus, understanding the principles governing condensate composition is critical to characterizing condensate function. The molecular bases of macromolecular composition have been studied in detail for several condensates^6-9^, but partitioning of small molecules into condensates remains poorly understood. Using mass spectrometry with validation by fluorescence microscopy, we quantified partitioning of ∼1700 metabolites and FDA-approved drugs into four condensates composed of different DNA and/or protein scaffolds. We found that partitioning spanned from ∼100-fold exclusion to ∼10,000-fold enrichment. Strong correlations between the different condensates suggest an underlying physical similarity despite disparate macromolecular components. Strongly partitioning molecules generally did not bind condensate-forming proteins with high affinity under conditions where condensates do not form, suggesting only a minor role for stereospecific interactions in partitioning. We developed a machine learning model that accurately predicts partitioning using only physicochemical properties of the compounds. The strongest predictors of partitioning were features related to aqueous solubility and hydrophobicity. Small molecule partitioning was similar for condensates in aqueous buffer and in concentrated cell extracts, suggesting analogous behaviors in vivo. Together, the data and model suggest that small molecule partitioning is not generally based on high-affinity, stereospecific interactions with scaffolds, but rather on physical properties of the compounds, and their compatibility with an emergent hydrophobic environment within the condensate. Our results will aid design of chemical compounds that target biomolecular condensates and reveal unexpected physical and functional similarities of distinct condensates.

## Introduction

Biomolecular condensates are cellular compartments that concentrate proteins, nucleic acids, and likely small molecules in the absence of an encapsulating membrane^1,2^. Many condensates appear to form through liquid-liquid phase separation of multivalent macromolecules^1,2,10^. Condensates are involved in numerous biological processes, including signal transduction, intermediary metabolism, RNA metabolism and gene regulation. Aberrant condensates are associated with human diseases including neurodegeneration, cancer and viral infections^5,11,12^, and bioinformatic analyses have recently implicated condensate dysregulation in over 1000 genetic disorders ^13^. Condensates have thus emerged as therapeutic targets ^12,14,15^. Drug development efforts seek to identify small molecules that disrupt condensates or modulate their material properties, or that alter the activity of specific macromolecular components.

Condensates are believed to segregate, concentrate, and modulate biochemical processes in vivo, affording specificity and efficiency to individual reactions and pathways^3-5^. Understanding the mechanisms that determine condensate composition is thus important for defining condensate functions. In models of macromolecular composition, many proteins and RNAs are recruited through specific binding to other condensate components^6-9^. Additionally, proteins can be enriched in or excluded from condensates without apparent site-specific binding, in a manner depending on a protein’s overall charge^16-18^. Much less is understood about the factors that govern the composition of small organic molecules in condensates^19^. Multiple mechanisms are likely to be important and can be conceptualized on a spectrum of binding affinities and specificities. At one end, high affinity, stereospecifically-defined interactions can occur between a compound and complementary element in a macromolecular condensate component. These will drive recruitment of the compound into the condensate, as has been reported for enrichment of certain anti-cancer drugs in condensates containing their targets^20^. In contrast, low affinity, low specificity interactions with solvent and/or other molecules can govern the distribution of a compound between the distinct biochemical milieus inside and outside a condensate. Such factors can lead to enrichment or exclusion of a compound from the condensate phase. It remains unknown which types of interactions play dominant roles in dictating the small molecule recruitment into biomolecular condensates.

Here we examined the partitioning of a large collection of small molecule metabolites and drugs into four different condensates composed of unrelated proteins and protein-DNA mixtures, both purified and in cell extracts. The degree of partitioning spans nearly six orders of magnitude, from ∼100-fold exclusion to ∼10,000-fold enrichment. Surprisingly, partitioning is correlated among the different condensates, indicating a similarity in their underlying physical properties. Several highly partitioning compounds do not show measurable interactions with one model protein system in non-phase separating conditions, suggesting that enrichment is not due to specific binding to condensate components. We developed a machine learning model whose only inputs are physicochemical properties of the compounds, not molecular structures, that predicts partition coefficients with quantitative accuracy in both the pure protein and extract systems, providing potential routes to drugs that target condensates. Furthermore, the model indicates that the strongest predictors of partitioning behavior are aqueous solubility and related measures of hydrophobicity. The experimental data and modeling support that the partitioning of small molecules into condensates is defined by their chemical properties, wherein compounds with enhanced hydrophobic character favor concentration into the condensates. This enrichment of organic compounds in the condensate is an emergent biochemical property of the phase separated compartment, which is not found in the individual macromolecules.

### Partitioning of chemical compounds varies over nearly six orders of magnitude, and is correlated between condensates

We examined the partitioning of small molecules into four disparate condensates composed of proteins and nucleic acids. We characterized two synthetic condensates, one composed of polySH3 and polyProline-Rich-Motif (PRM)^21^, and one composed of polySUMO and polySUMO-Interaction-Motif (SIM)^6^. Additionally, we examined two condensates formed by naturally occurring macromolecules, one formed by the yeast P-body protein Dhh1^7^ and the other formed by the innate immune signaling protein cyclic GMP-AMP synthase (cGAS) and double stranded DNA^22^.

For each condensate, we assessed partitioning of two libraries of molecules, an ∼200 compound metabolite library, and an ∼1500 compound library of FDA approved drugs. To visualize the diversity of the compounds, we calculated ∼50 absorption, distribution, metabolic, and excretion (ADME) and chemical properties using QikProp^23^. The multidimensional descriptor set was embedded into a two-dimensional chemical space using Uniform Manifold Approximation and Projection (UMAP)^24^. Clustering analysis using an unsupervised learning algorithm (HDBSCAN)^25^ identified 10 groups of small molecules with similar chemical attributes, suggesting QikProp descriptors convey enough chemical information to discriminate between classes of molecules (Figure 1A and interactive Extended Data Figure 1). There are well-defined, isolated clusters for steroidal compounds, guanidine-containing compounds, and β-lactam antibiotics. The other clusters share boundaries but have distinct characteristics. The largest of these clusters contains small biomolecules, multi-chlorinated-aryl compounds, and common nonsteroidal anti-inflammatory drugs with a variety of structures. Overall, the UMAP and clustering indicate a relatively diverse, but sparsely populated set of molecules.

**Figure 1.**
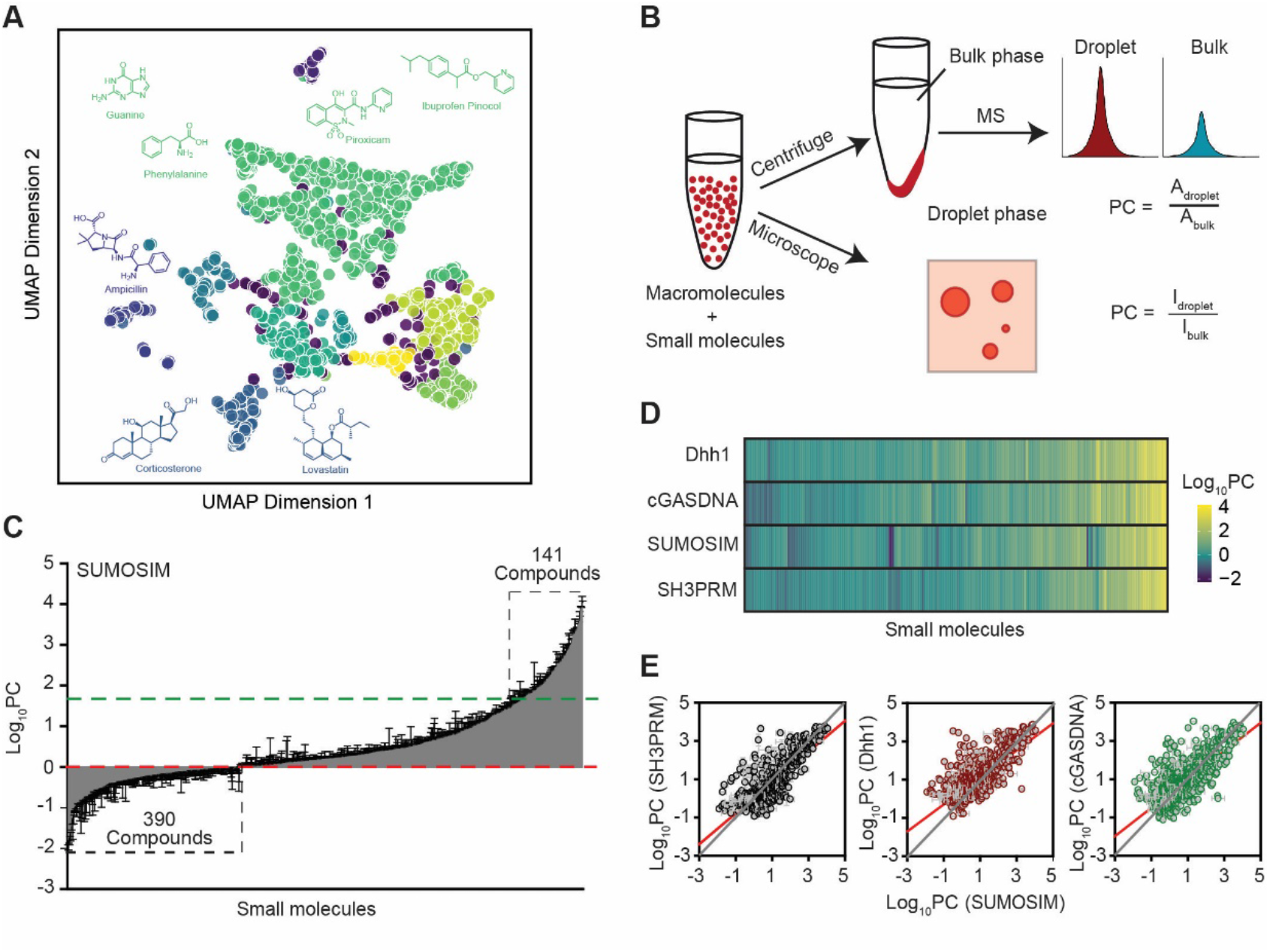
Partitioning of chemical compounds varies over nearly six orders of magnitude, and is correlated between condensates. A. UMAP representation of ∼1700 small molecules used in the analysis based on physical features generated in QikProp. B. Schematics illustrating assays to measure small molecule partitioning into condensates based on mass spectrometry (top) and confocal fluorescence microscopy (bottom). C. Bar chart of partition coefficients, ordered from smallest to largest, for 1216 compounds into SUMOSIM condensate. Red and green dotted lines indicate logPC = 0 and logPC_SUMOSIM_ (1.77), respectively. Number of compounds with logPC < 0 and logPC > logPC_SUMOSIM_ are indicated. D. Heat map showing logPC values for each of the four condensates indicated. E. Scatter plots of logPC values for compounds into SUMOSIM condensates versus into SH3PRM, Dhh1 and cGASDNA condensates. Error bars in panels C and D indicate standard error for two independent replicates.

We developed two assays to measure partitioning of the compound libraries (Figure 1B). In the first, we incubated sub-libraries of ∼300 compounds, each at 1 *µ*M concentration, with the macromolecules under phase separating conditions (2-10 *µ*M concentrations). After equilibration (1 to 24 hours), the condensates were collected by centrifugation followed by manual separation of pellet and supernatant. Compounds were subsequently extracted from the two samples and quantified by mass spectrometry, yielding the partition coefficient (PC) as the ratio of concentrations in the pellet and supernatant. Our pooled compound library approach relies on each compound behaving independently, with compounds not causing significant exclusion or recruitment of other compounds into condensates. To examine this, we analyzed partitioning into SUMOSIM condensates of a 240-compound group from the drug library and the same molecules divided into three 80-compound sub-groups. Apart from a few outliers (∼5 molecules), the partition coefficients measured in each case were similar (R^2^ =0.86) (Extended Data Figure 2). Thus, while we cannot entirely rule out crosstalk between certain molecules, the compounds generally behave independently from each other in this assay.

To validate the mass spectrometry results, we used confocal fluorescence microscopy where the 30 fluorescent molecules in the drug library were individually incubated with macromolecules under phase separating conditions. PCs were determined from the ratio of fluorescence intensity within the condensate and surrounding solution. In each of the condensates, partition coefficients measured by mass spectrometry and microscopy were correlated (R^2^ = 0.55 – 0.75), supporting the accuracy of both approaches (Extended Data Figure 3). The modest number of outliers was likely due to changes in fluorescence intensity of certain fluorescent compounds in the dense and dilute phases^26^, or crosstalk between molecules in the mass spectrometry assay.

Figure 1C plots the PC values of the metabolite and drug libraries in the SUMOSIM condensate measured using the mass spectrometry assay. The values span a nearly 1,000,000-fold range, from ∼0.01 to 10,000, with the strongest partitioning corresponding to a free energy of ∼5.6 kcal/mol for transfer of a compound into the condensate from the diluted phase. Of the 1216 compounds that could be reliably measured (see Methods), 215 had PC values < 0.8, indicating that they are substantially excluded from the condensates, 330 compounds had PC values in the range 0.8-1.5, indicating little preference for recruitment into condensates over the surrounding solution, and 660 had PC values of 1.5 – 10,000 indicated modest to strong enrichment in the condensate. Given the droplet volume fraction of 1.2 ± 0.2 % and 1 *µ*M total compound concentration, the highest enrichment corresponds to ∼90 *µ*M concentration in the condensate. The metabolites are generally limited to the lower end of the PC range, spanning 0.14 – 35, while the drugs sample the entire range of values.

We initially anticipated that the individual condensate systems would show different distributions of partitioning since they are composed of unrelated molecules with different physical properties (Table S6). Surprisingly, however, the different condensates show similar ranges of PC values for the small molecules (Figure 1D and Extended Data Figure 4) and the distributions are strongly correlated between the different condensates over the full range of PC values (Figure 1E, Extended Data Figure 5). Correlations are weaker when considering smaller ranges of PC values (less than 1∼2 log units) (Extended Data Figure 6) and may indicate that effects specific to each macromolecule govern small differences in PC. These results indicate that despite the differences in macromolecular components there is an underlying physical similarity between the condensates such that the same small molecules tend to be recruited or excluded in each condensate system.

### Partitioning is not governed by stereospecific binding to condensate scaffolds

Several features of our PC dataset suggest that partitioning of the small molecules is not governed by specific binding to the condensate macromolecular components. First, for enrichment due to stereospecific binding to a discrete site, PC_compound_ is bounded by 1 and PC_scaffold_, assuming the macromolecular scaffolds have equal affinity for a compound in the dense and dilute phases^6^. Yet here, each condensate has numerous compounds that are substantially excluded (PC<1), and compounds with PC values that far exceed those of the macromolecular components (Figures 1C, D, Extended Data Figure 4). Second, while PC_compound_ values can lie outside of these limits if the macromolecular components have different affinities for compounds inside and outside of the condensate^27^, the similarity of distributions between the different condensates argues against this possibility (Figure 1E, Extended Data Figure 5); it is unlikely that two independent scaffolds would both possess stereospecific binding sites for a compound that are differentially populated to similar degrees in the dense and dilute phases. Finally, the highly partitioning compounds for each condensate do not possess common structural features, as would be expected if they recognized a specific binding site in the scaffold (Extended Data Figure 7). Thus, the data suggest that partitioning is not driven by stereospecific binding to condensate scaffolds, but rather by general physical properties of the compounds, which differentially favor the dense or diluted phase.

To test this hypothesis, we used isothermal titration calorimetry (ITC) to examine binding of eight compounds to the polySUMO-polySIM complex at a concentration below the phase separation threshold (20 *µ*M module concentration). These compounds had PC values ranging from 50 to 800, and were titrated up to 200 *µ*M, a 10-fold excess over the scaffolds. As illustrated in Figure 2 and Extended Data Figure 8, at both 25 °C and 35 °C, four of the compounds showed flat or nearly flat titration profiles, with enthalpy of binding near zero. The absence of measurable heat at two different temperatures strongly suggests the absence of binding, rather than a small heat of binding. Two of the compounds showed flat profiles with non-zero enthalpies, suggesting a weak interaction that is far from saturation even at 10-fold excess ligand. Only two of the compounds showed saturable binding, both with *K*_D_ values in the low micromolar range. The binding behavior does not appear to be related to partitioning, as the two compounds with micromolar affinity are intermediate in the PC range sampled (Extended Data Figure 8). Thus, most of the compounds examined do not have measurable affinity for the polySUMO-polySIM complex in the absence of phase separation. These experiments further support the hypothesis that partitioning is driven primarily by the physical properties of compounds and condensates rather than stereospecific binding to scaffolds.

**Figure 2.**
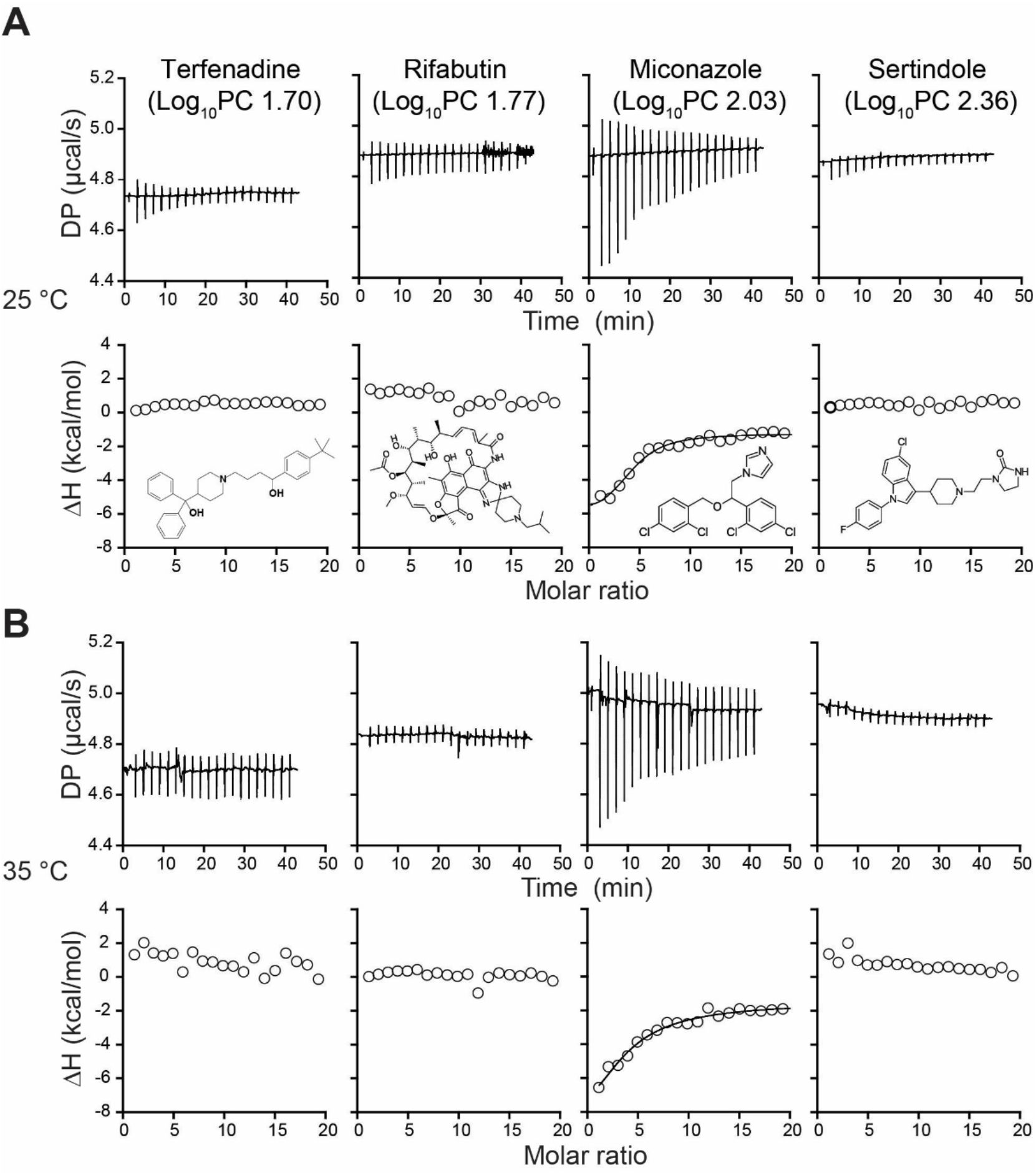
Many compounds do not bind SUMOSIM scaffolds under non-phase separating conditions. Raw thermograms (top row) and integrated enthalpies (bottom row) of titrations of terfenadine, rifabutin, miconazole and sertindole (left to right) into 20 *µ*M module concentrations of polySUMO + polySIM (below phase separation threshold) monitored by ITC. A and B show titrations at 25 °C and 35 °C, respectively.

### A predictive machine learning model for partitioning based on compound physical properties

We turned to statistical modeling to determine which chemical features of small molecules determine partitioning, with a related goal of exploiting the models to predict the partitioning behavior of new compounds. We used the chemical space map and clustering to define balanced training and test sets for modeling and validation purposes. We withheld 20% of the molecules from each cluster for external validation. Of the remaining molecules, 70% were randomly selected as the training set to develop models, and 30% were used as a test set to compare modeling strategies. This included evaluation of different data structures and ML algorithms through comparison of the test set R^2^ and mean absolute error (MAE) in predicted logPC of the resultant models (see Supplementary Materials for modeling details).

Extreme gradient parallel tree boosting (XGBoost) models generally outperformed other algorithms^28^, and QikProp descriptors outperformed structure-based molecular encodings that produce a larger set of features (Table S7 and Supporting Information). Adding descriptors of the condensate scaffolds (Table S6) led to overfitting and these features were not used in our final models. The XGBoost model of SUMOSIM partitioning had a modest test R^2^ of 0.47 and a logPC MAE of 0.49 (Figure 3A). Models for the other condensate systems performed similarly, with test R^2^ ranging 0.48-0.51 and logPC MAE 0.49-0.43 (Extended Data Figure 9).

**Figure 3.**
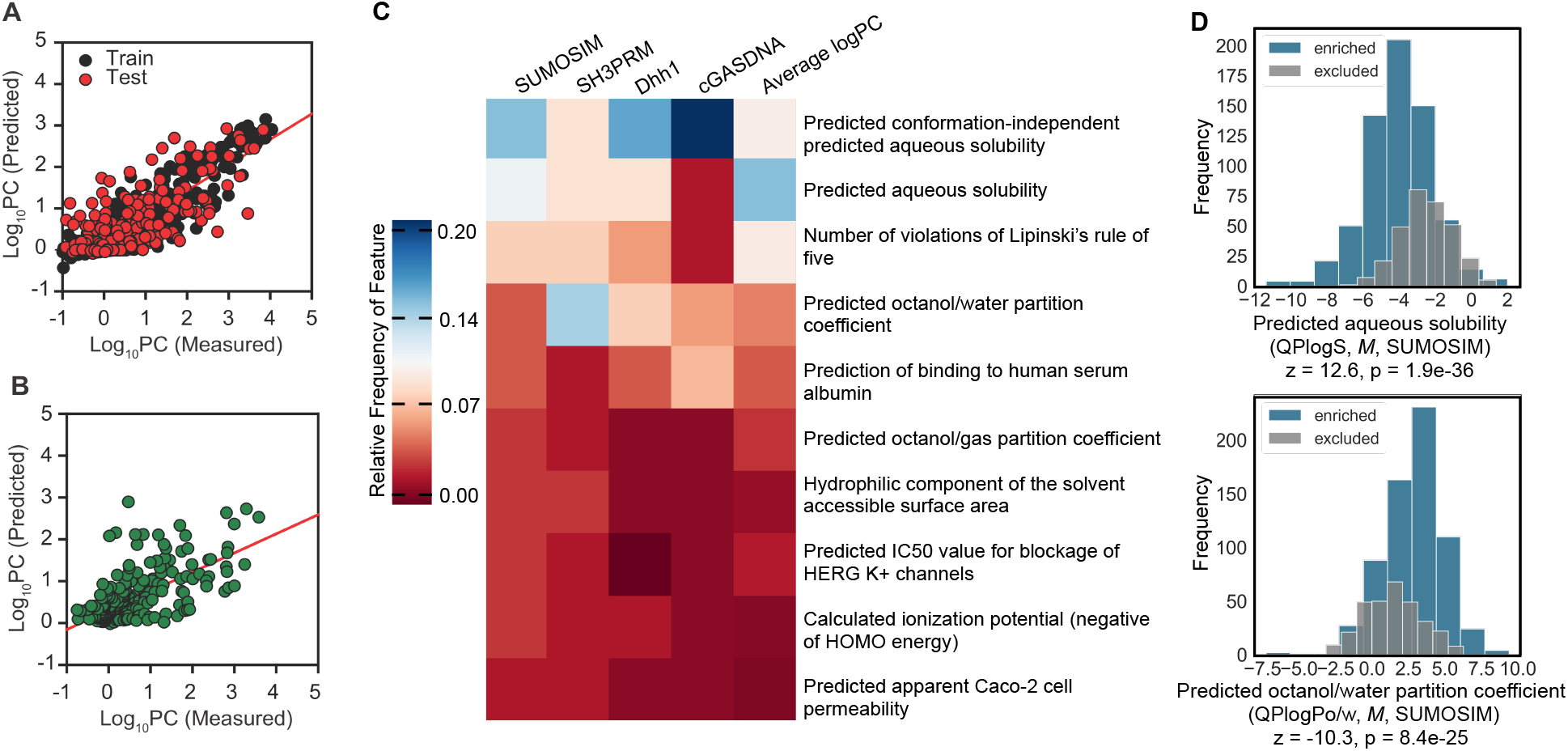
Machine learning models for partitioning of small molecules into biomolecular condensates are predictive and indicates physical features determine partitioning behavior. A. XGBoost model of small molecules partitioning into buffered SUMOSIM condensates (N train = 656 (black), N test = 302 (red), test R^2^ = 0.47, test MAE = 0.494). B. Validation set predictions for the average logPC model; (N train = 1145 (not shown), N validation = 263 (green), validation R^2^ = 0.42, validation MAE = 0.44). C. Heatmap showing the relative feature importance of the top 10 features in the individual condensate and average logPC models. D. Histogram comparison of the distributions model features for molecules that are enriched (N = 651, blue) vs. excluded (N = 307, gray) from SUMOSIM condensates. *Z*-test analysis indicated that the mean predicted aqueous solubility of enriched molecules is lower than excluded molecules and that the mean predicted octanol/water coefficient is higher for enriched molecules.

Given the strong correlation of PC values across the condensate systems, we reasoned that performance of our models should not depend strongly on which condensate PC dataset is used for model training. To address this, we compared the individual models to one based on the combined data for all four condensates (Table S8). This combined model had improved R^2^ and MAE, of 0.51 and 0.46, respectively. To assess whether this improvement resulted from the larger training sample size, we also modeled the average logPC for each molecule across all condensates, a data size comparable to the individual condensate models (Table S8, Entry 6). The average logPC model had a test R^2^ of 0.50 and the overall best MAE, 0.41. Thus, combining the data from the four condensates, rather than a larger sample size per se, improved the model predictions. The better performance of the combined and average logPC models suggests that condensate-small molecule interactions are conserved with respect to condensate identity. This emergent quality implies that condensate composition plays a secondary role in small molecule partitioning.

We validated the average logPC model by predicting the withheld data. The model successfully predicted partitioning behavior for the validation set within one-half of a logPC unit (validation R^2^ = 0.42, MAE = 0.44, Figure 3B), corresponding to PC error of < 3-fold. Although the validation predictions plateau near logPC = 0 (likely due to the limited experimental range of logPC for excluded molecules compared to enriched molecules), the MAE of the excluded validation points is only 0.34 logPC units. We recognize that the sparsity of data relative to the molecular diversity surveyed resulted in the observed undertraining and aim to supplement our database of small molecule partitioning measurements to explore this limitation in future studies. Prediction errors did not substantially change as a function of which cluster a compound originated from, suggesting that the model effectively covers the diverse compound representation. Furthermore, only 12% of molecules had prediction errors greater than one logPC, showing that our model is effective in differentiating highly enriched molecules from those that are weakly attracted to or excluded from the condensates. We anticipate the ability to predict differences in partitioning across nearly six orders of magnitude will have significant benefit for future applications.

An advantage of tree-based algorithms like XGBoost is that they afford insight into the relative importance of the QikProp features. In this regard, descriptors of solubility/hydrophobicity dominated the SUMOSIM and average logPC models (Figure 3C and Extended Data Figure 10). *Z*-test comparisons of the top features in the SUMOSIM model demonstrated a significant difference in the distributions of enriched vs. excluded small molecules (Figure 3D, Extended Data Figure 11). Furthermore, the conservation of feature importance across models of the four individual condensates suggests that the rules of small molecule partitioning are relatively consistent for all condensates (Extended Data Figure 10). These observations provide further support for the hypothesis that partitioning is largely driven by the physical properties of the compounds, as opposed to stereospecific binding.

### Solution conditions can significantly influence small molecule partitioning

Our experimental and modeling results suggest that the chemical environment within a condensate is different from the surrounding solution, and this difference plays an important role in determining whether organic compounds are enriched or excluded. However, the vastly more complex environment of the cell relative to simple buffers might impact partitioning in ways not accounted for by our experiments thus far. To address this, we quantified partitioning of the drug library into SUMOSIM condensates under conditions designed to mimic the cellular environment, either in a relatively dilute U2OS cell lysate (∼4 mg/ml total protein) or a highly concentrated *Xenopus laevis* oocyte extract (80-100 mg/ml, comparable to the eukaryotic cytoplasm). As shown in Figure 4A,B, for both systems the partitioning behaviors remain modestly correlated to those observed for SUMOSIM condensates in simple buffer (R^2^ of 0.39, 0.34, respectively). However, the total range of PC values decreases by ∼100-fold, spanning 0.1 ∼ 1000, with only a few values larger than 100. The smaller range is likely due in part to an ∼10-fold decrease in PC values for the polySUMO and polySIM proteins themselves (60 in buffer, 8 in U2OS lysate, 5 in *Xenopus* extract). Additionally, the much greater complexity of the lysate/extract solution likely dampens differences between dense and dilute phases. On average, the logPC values decrease ∼2-fold between buffer and either lysate, as indicated by the slopes of the lines correlating the distributions, and the relatively large root-mean-square-deviation (RMSD) in logPC values between the datasets (1.04 and 0.97, respectively, for buffer vs lysate and buffer vs extract). Although the lysate and extract have disparate total protein concentrations, their effects on compound partitioning are similar, with R^2^ of 0.47 and RMSD of 0.45 logPC units between the datasets (Figure 4C). Thus, the differences between the buffer and lysate/extract environments appear to stem more from the number of molecular species rather than the concentrations of those species.

**Figure 4.**
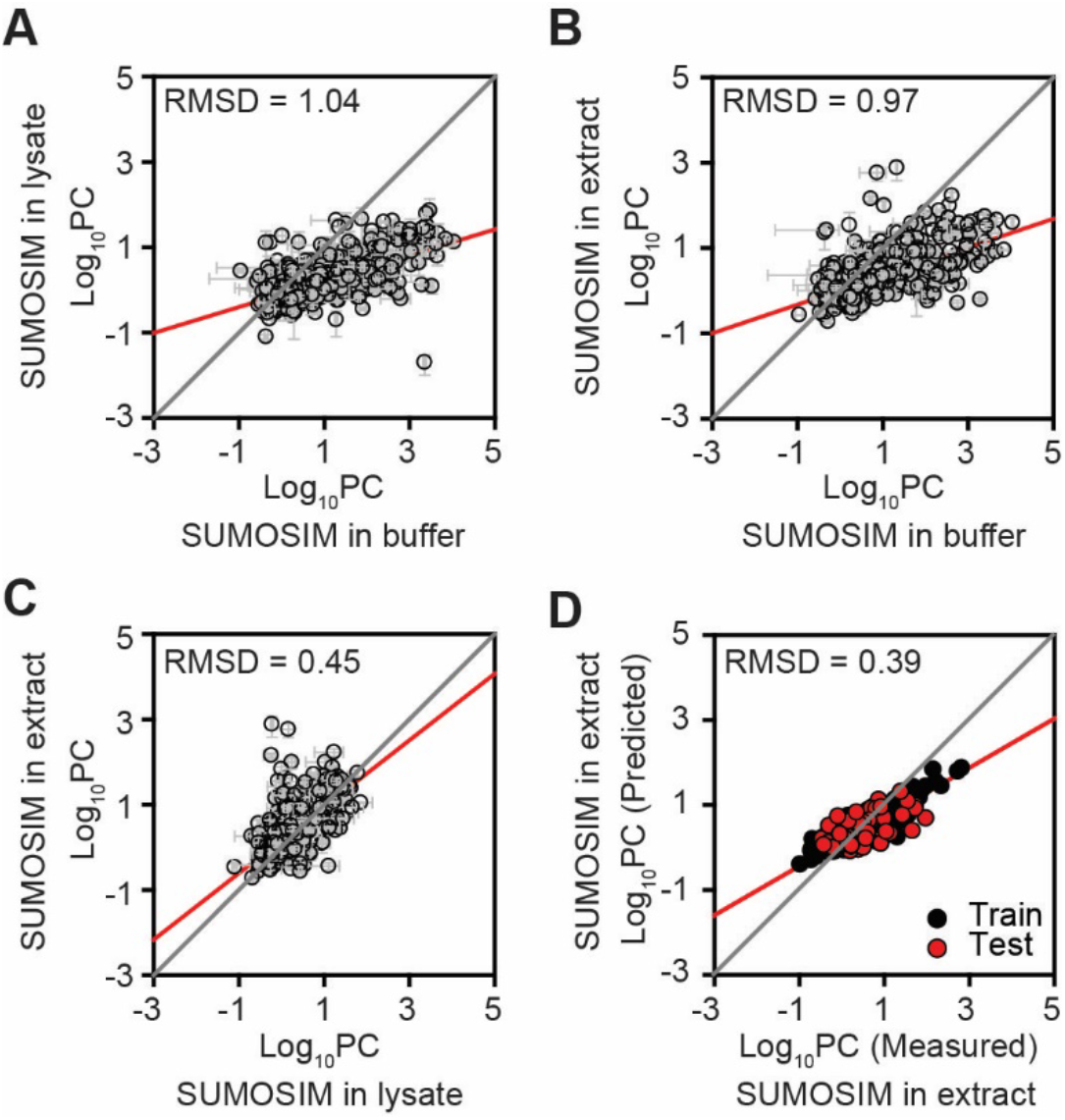
Compounds show different distributions of PC values in different solvent conditions. A, B. Scatter plots of logPC values of compounds into SUMOSIM condensates generated in U2OS cell lysates (A) or in Xenopus oocyte extracts (B) vs in buffer. C. Scatter plot of logPC values of compounds into SUMOSIM condensates generated in Xenopus oocyte extracts versus in U2OS cell lysates. D. Machine learning model of compound partitioning into Xenopus oocyte extracts. Error bars indicate standard error for two independent replicates.

As with the pure condensate systems, we were able to generate a features-based statistical model of compound partitioning into the SUMOSIM condensates in the *Xenopus* extract system (Figure 4D). The model was somewhat less predictive in the more complex solution than in simple buffer (validation R^2^ = 0.39, validation MAE = 0.30), although this deficit may be due to the reduced number of compounds that could be analyzed due to technical limitations (see Supporting Information) rather than an inherent limitation of modeling extract systems. The features driving the model in the extract system shared similarities with the buffer model; hydrophobicity effects still dominate feature importance, although measures of solubility are not important in the extract model (Extended Data Figure 12). Together the data and modeling demonstrate that even in highly complex solution environments mimicking the cytoplasm, the drugs show a wide range of PC values that can be predicted by their physical properties.

## Discussion

We have demonstrated that the partitioning of small molecules into condensates varies over nearly six orders of magnitude, from modest exclusion to strong enrichment. The combination of biochemical data and statistical modeling indicates that in most cases partitioning is largely driven by the hydrophobicity and aqueous solubility of compounds, rather than their ability to stereospecifically bind to discrete sites on condensate scaffolds. An unexpected similarity of partitioning profiles among several different condensates suggests that phase separation results in a shared, general physical or chemical property among the macromolecules we have characterized. This property differentiates the physical environment within condensates from that of the surrounding solution, leading to enrichment or exclusion. The physical basis of the difference in environment remains unclear but may involve solvent structure within condensates or perhaps a greater abundance of transient, partially unfolded states of proteins that could create hydrophobic surfaces to recruit small molecules non-stereospecifically^29,30^.

A variety of approaches have been suggested for modulating the activities and/or physical properties of condensates to impact human diseases ^3,12,14^. The machine learning methods we have deployed here could accelerate these approaches by facilitating rapid identification of highly-enriching compounds. In diseases where therapeutic targets reside in condensates, modeling could guide rational modification of existing inhibitors to increase their concentrations in the compartments. Existing drugs could also be grafted onto strongly partitioning compounds to direct them into condensates; both of these strategies could provide increased potency or selectivity toward targets. It is important to note that enrichment via site-specific binding of a small molecule to one condensate component does not imply a thermodynamic advantage for binding and modulating the properties of any other component. Thus, the potency of a drug targeting one condensate component might not improve simply by adding moieties that specifically bind another component. However, our finding that strong enrichment can be achieved without stereospecific binding, potentially due to solvent properties, suggests that drug potency could be enhanced by an overall increase in compound concentration within the condensate not localized at any specific binding sites.

Physical properties of condensates that form through phase separation, such as viscoelasticity and surface tension, are emergent. That is, they are inherently macroscopic, resulting from interactions of the parts of the system but not manifest in the parts individually^31^. Here we have found that the ability to recruit small molecules is an emergent biochemical property of SUMOSIM condensates, and likely others. Most drugs do not appear to bind the polySUMO-polySIM complex in the absence of phase separation, but are recruited strongly into SUMOSIM condensates. Thus, the free energy of partitioning emerges only upon phase separation. We also previously demonstrated that membrane dwell time, and consequently signaling activity, is higher in Nephrin/Nck/N-WASP condensates than in small complexes of the same molecules, again emerging as a function of size^32^. Future work on how biochemical activity resident in condensates varies as a function of condensate size will reveal the extent to which emergent biochemistry is a general feature of condensates in biology.

## Supporting information

Interactive_UMAP

## Figures and Figure Legends

The HTML file encoding Extended Data Figure 1, interactive_UMAP.html, will be included with the manuscript files (it cannot be embedded in a word document).

**Extended Data Figure 1**. Interactive chemical space map of all compounds used in this study. Hovering over data points shows the corresponding name and structure; molecules are colored according to their cluster.

**Extended Data Figure 2.**
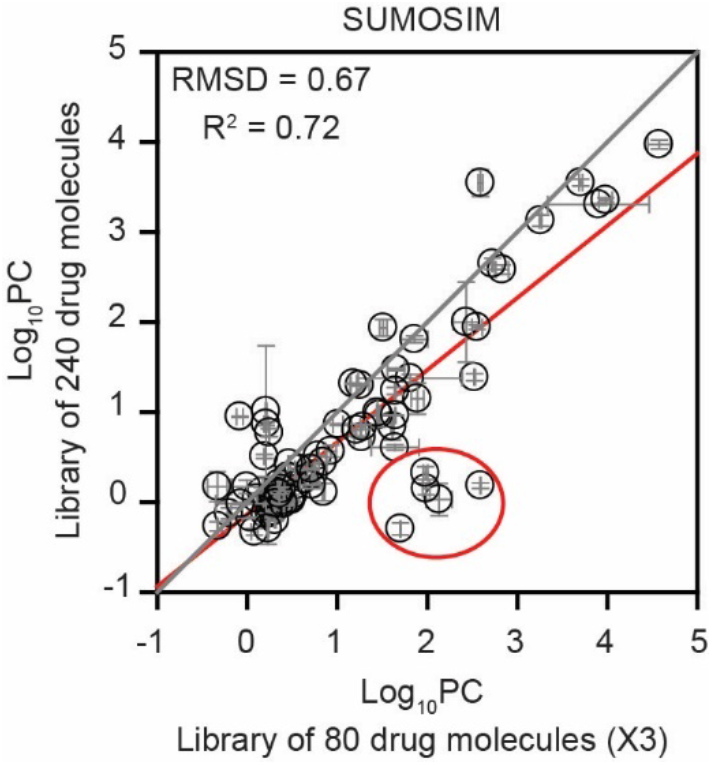
Most compounds have similar PC values when measured in large or smaller groups. Scatter plot of logPC values measured for a library of 240 compounds versus the values for the same compounds divided into three groups of 80 species. Other than the five compounds circled, most molecules have very similar logPC values in the larger or smaller groups. Error bars indicate standard error for two independent replicates.

**Extended Data Figure 3.**
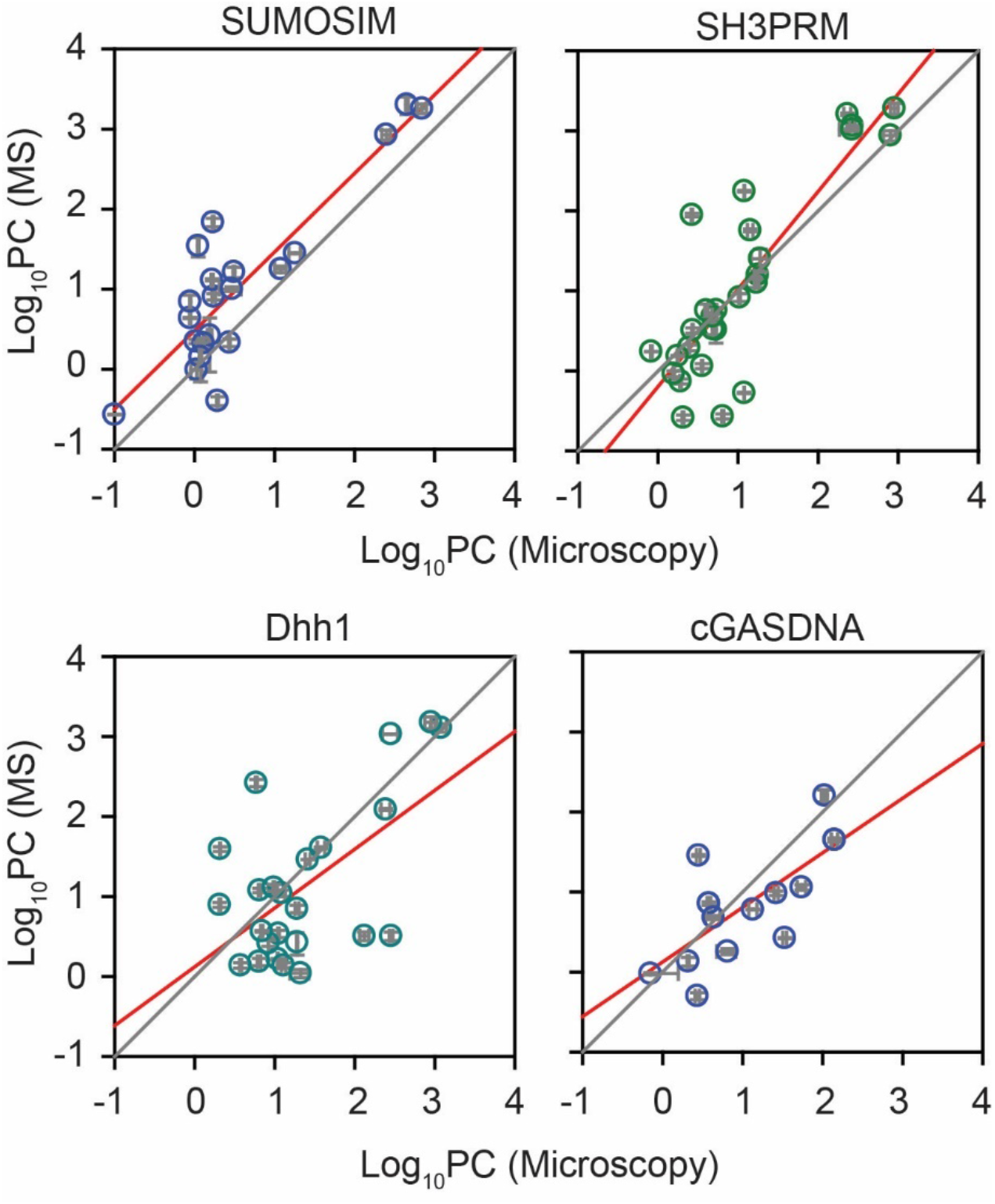
The mass spectrometry and confocal fluorescence microscopy assays of partition coefficients yield similar values for most species. Plots show correlations between logPC values measured by mass spectrometry and confocal fluorescence microscopy for the indicated condensates. Red and black lines show linear fit of the data and diagonal, respectively. Error bars indicate standard error for two independent replicates.

**Extended Data Figure 4.**
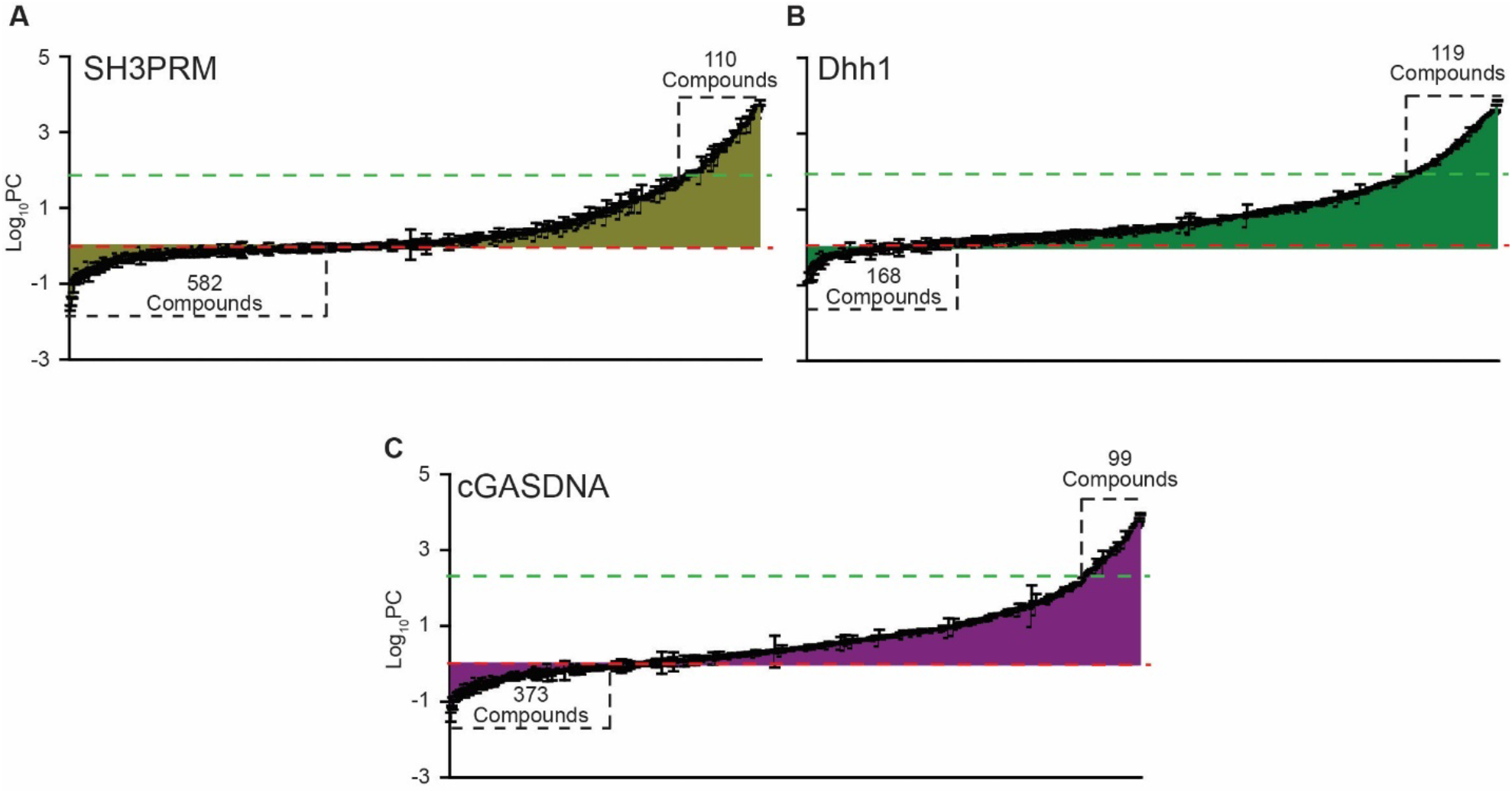
Each condensate shows a wide range of partition coefficients for the small molecules. Bar chart of partition coefficients, ordered from smallest to largest, for 1296, 1213, and 1296 compounds into SH3PRM (A), Dhh1 (B) and cGASDNA (C) condensates, respectively. In each panel the red dotted lines indicate logPC = 0 and green dotted lines indicate logPC_scaffold_ (2.0, 2.1, and 2.3, respectively). Error bars indicate standard error for two independent replicates.

**Extended Data Figure 5.**
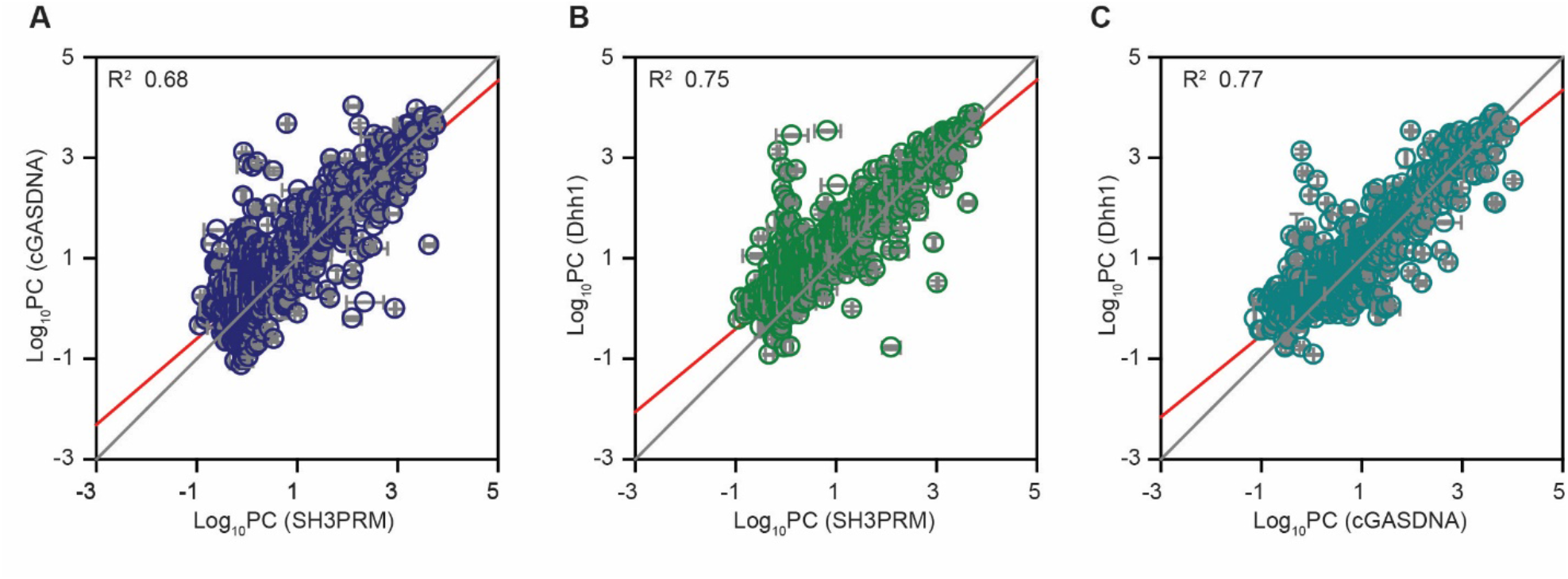
Partitioning of small molecules is correlated between the different condensates. Scatter plots of logPC values for compounds into A) cGASDNA versus SH3PRM condensates, B) Dhh1 versus SH3PRM condensates, and C) Dhh1 versus cGASDNA condensates. Error bars indicate standard error for two independent replicates.

**Extended Data Figure 6.**
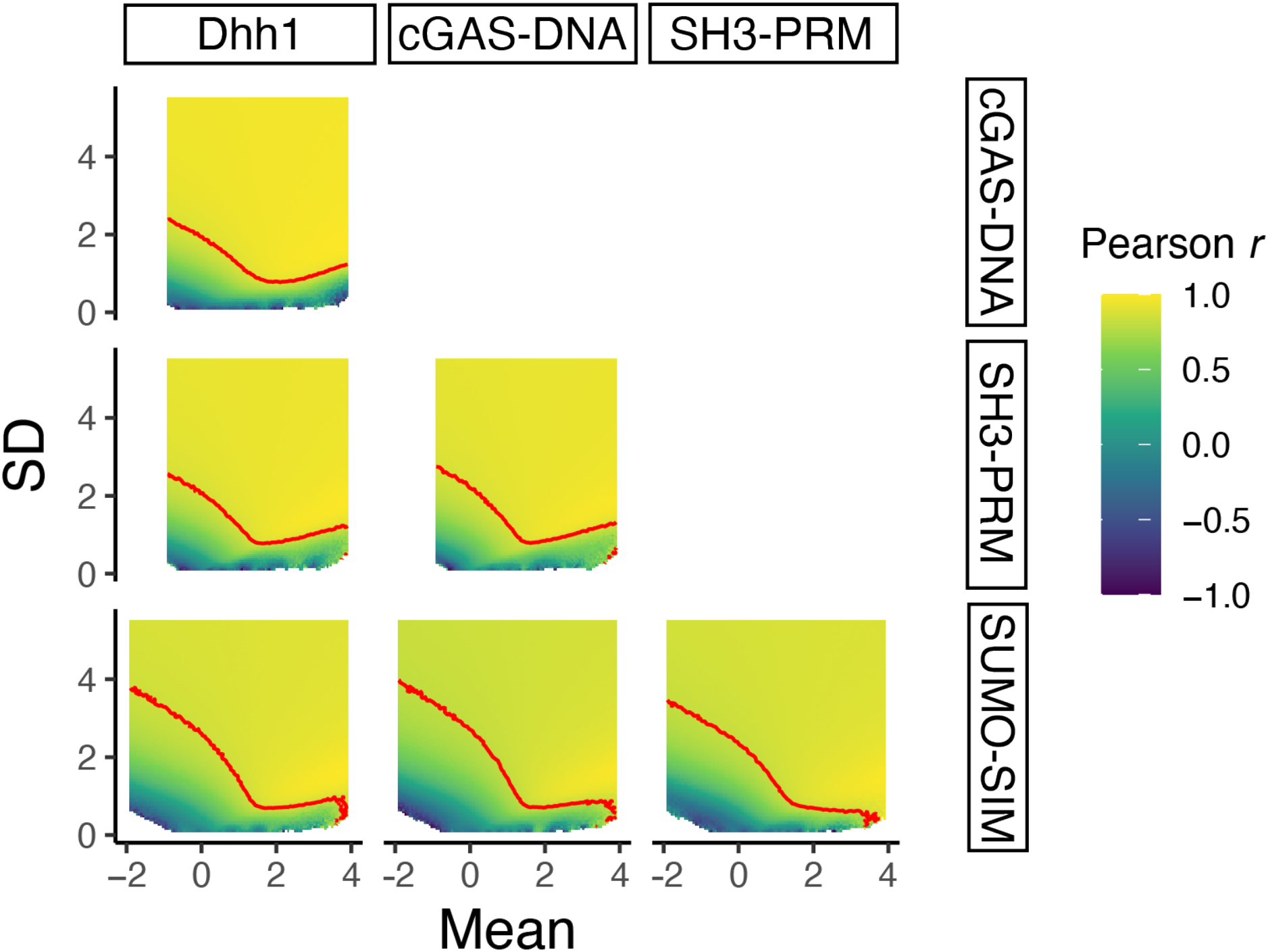
Correlations in subsampled windows between PCs of small molecules in different condensate systems. Each pixel in the raster images represents the Pearson correlation coefficient *r* between PC values of small molecules in each pair of condensate systems, with the PC data subsampled with a normally distributed bias with mean and SD parameters as indicated on the x- and y-axes, respectively. The red contour line indicates the *r* value for the complete (i.e., not subsampled) dataset. The images are thus a qualitative representation of the range of PC values over which correlations between each pair of condensate systems become apparent. The color scale is the same in all images.

**Extended Data Figure 7.**
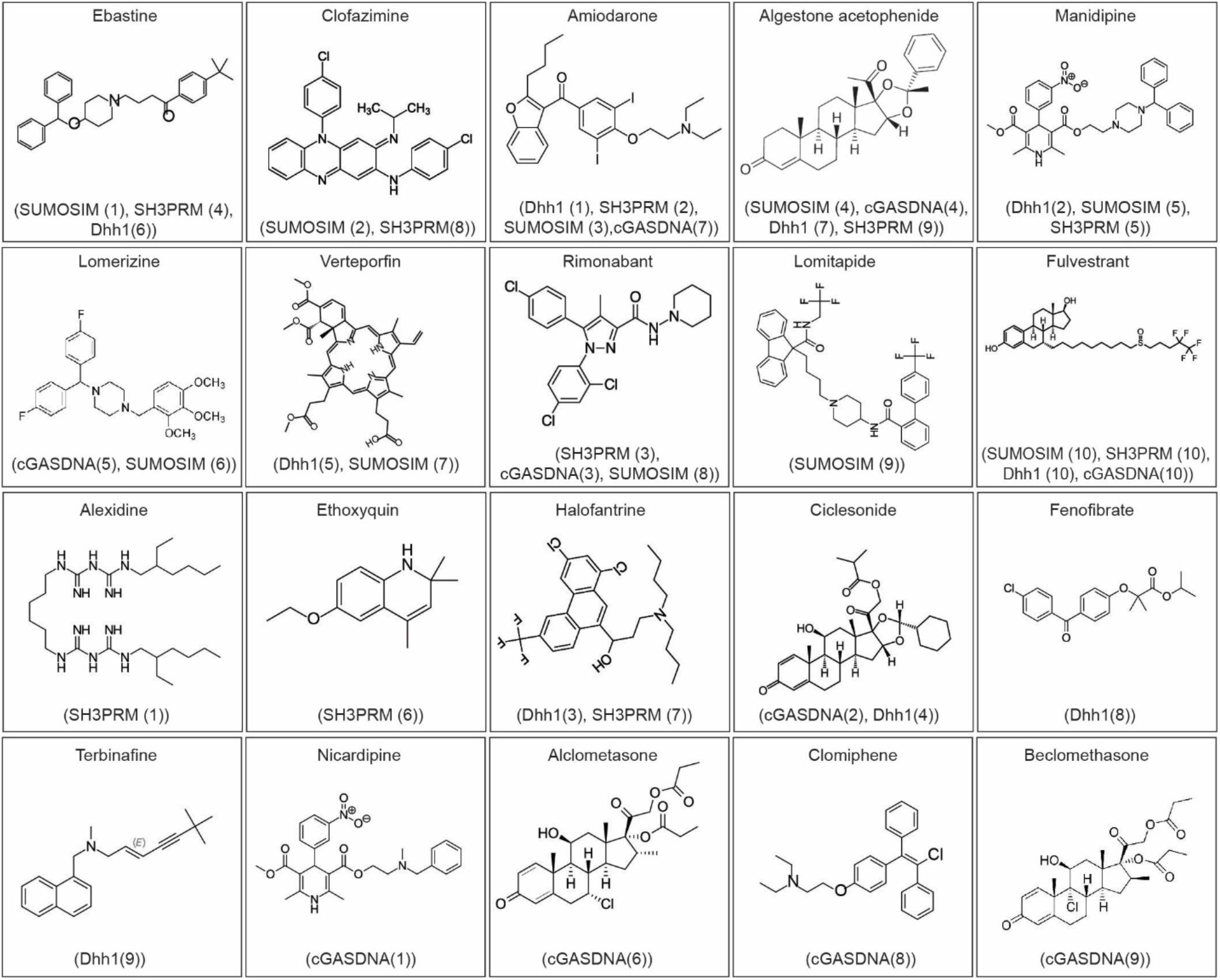
Chemical structures of the ten most strongly partitioning compounds for each of the four condensates.

**Extended Data Figure 8.**
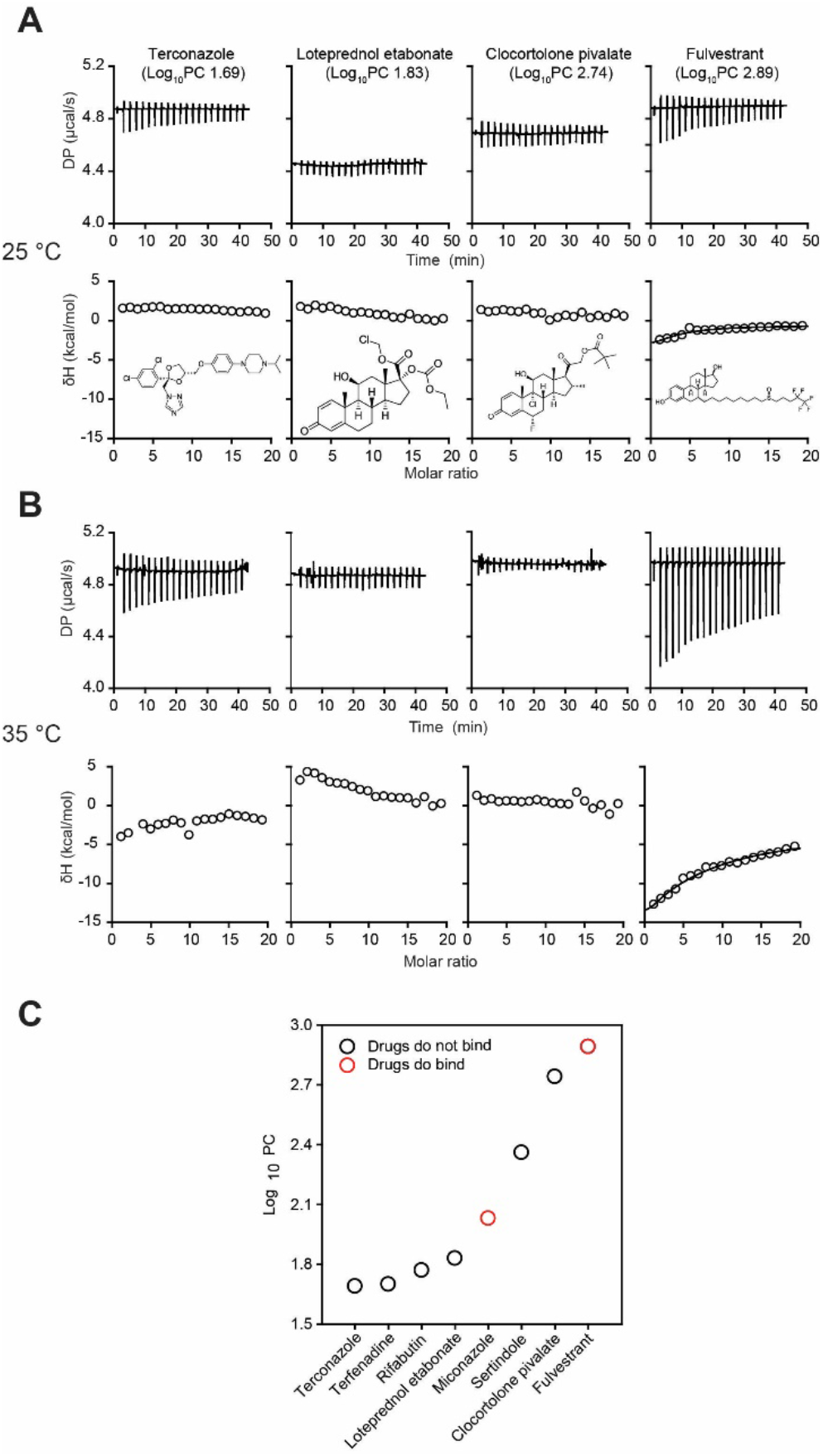
Additional compounds do not bind SUMOSIM scaffolds under non-phase separating conditions. Raw thermograms (top row) and integrated enthalpies (bottom row) of titrations of terconazole, loteprednol etabonate, clocortolone pivalate, and fulvestrant (left to right) into 20 *µ*M module concentrations of polySUMO + polySIM (below phase separation threshold) monitored by ITC. A) and B) show titrations at 25 °C and 35 °C, respectively. C) Summary of ITC data plotted against logPC for the eight compounds examined. Only miconazole and fulvestrant show saturable binding (colored red).

**Extended Data Figure 9.**
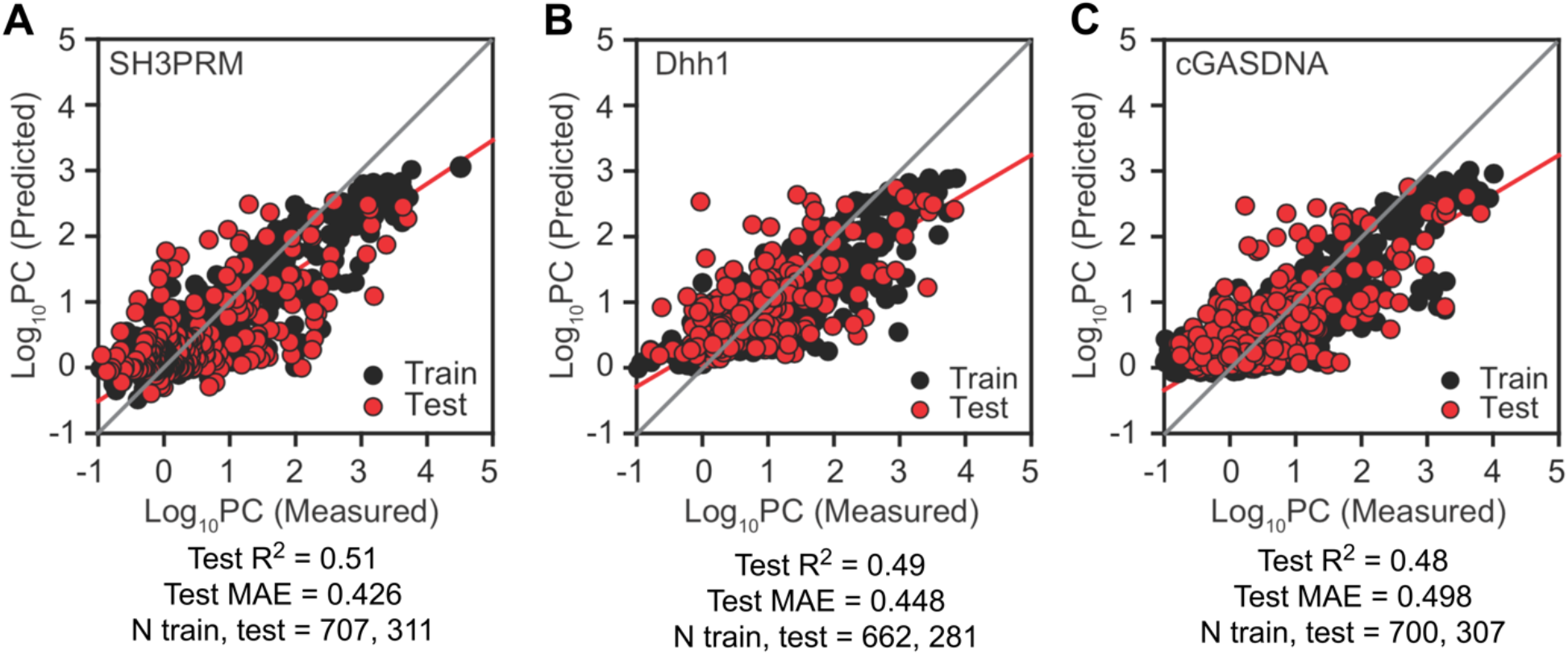
Extreme gradient boosting (XGBoost) models of other condensates. Shown are models of A) SH3PRM, B) Dhh1, and C) cGASDNA where the red and black lines show linear fit of the data and diagonal, respectively. MAE = mean absolute error.

**Extended Data Figure 10.**
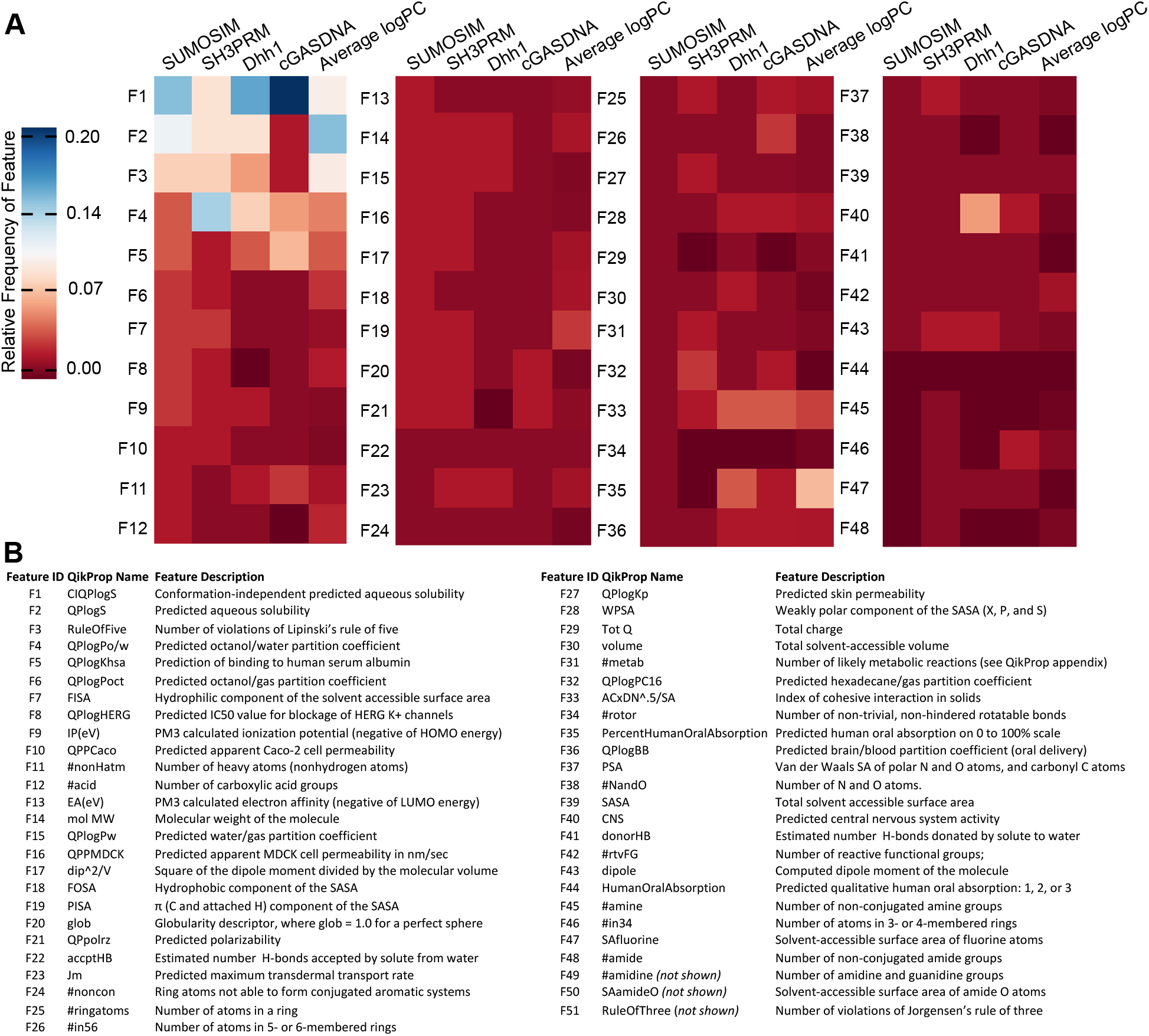
Heat map of the molecular descriptor importance for the four condensates. A. Feature importance is computed as the relative frequency by which a feature appears in the ensemble of trees. B. Feature identifiers with the corresponding QikProp names and descriptions.

**Extended Data Figure 11.**
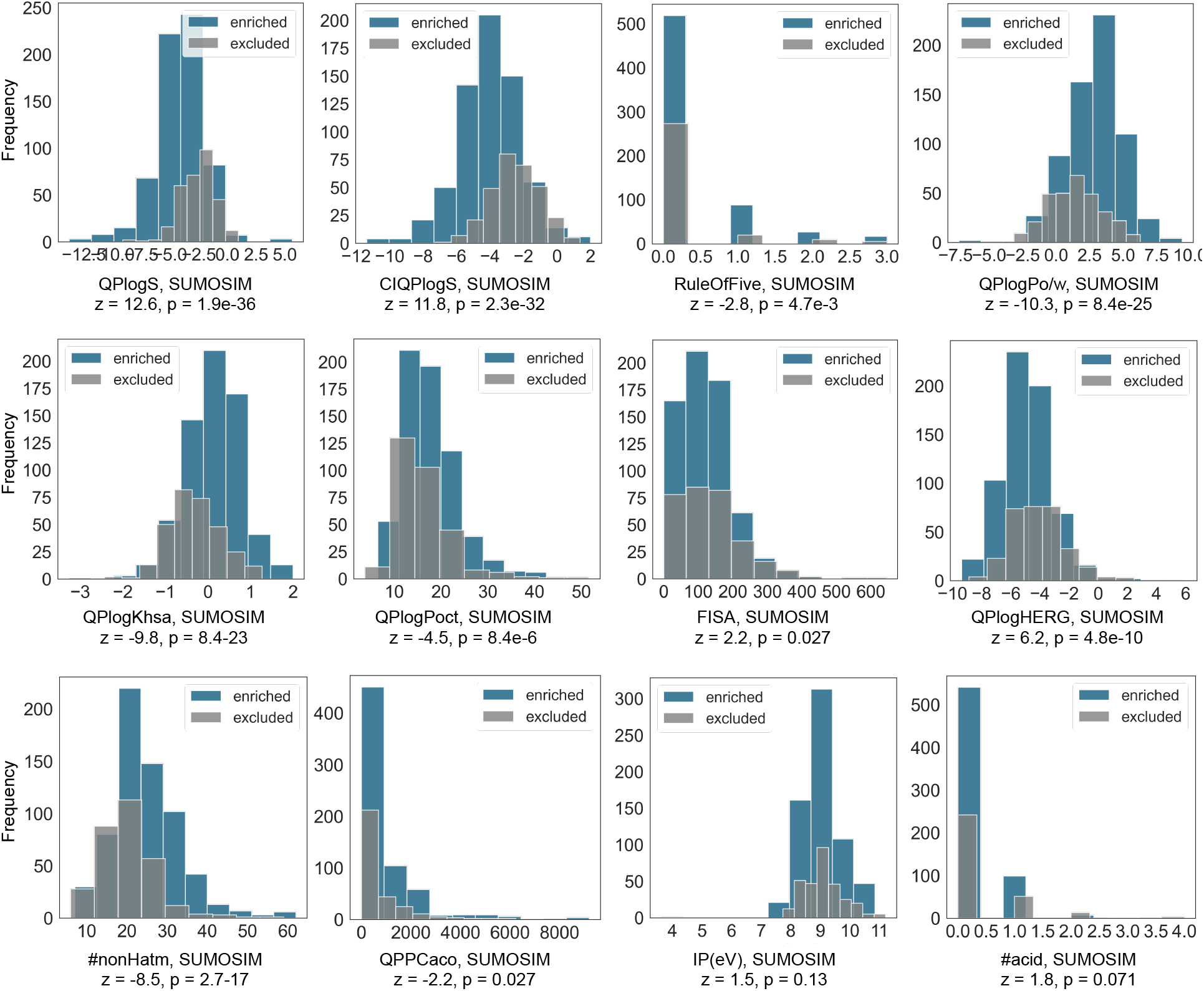
*Z*-test comparison of enriched (N = 651) vs. excluded (N = 307) molecules for the top features in SUMOSIM model. Enriched molecules (blue) had measured logPC greater than zero, and excluded molecules (gray) had measured logPC less than or equal to zero. Feature descriptions are provided in Extended Data Figure 11.

**Extended Data Figure 12.**
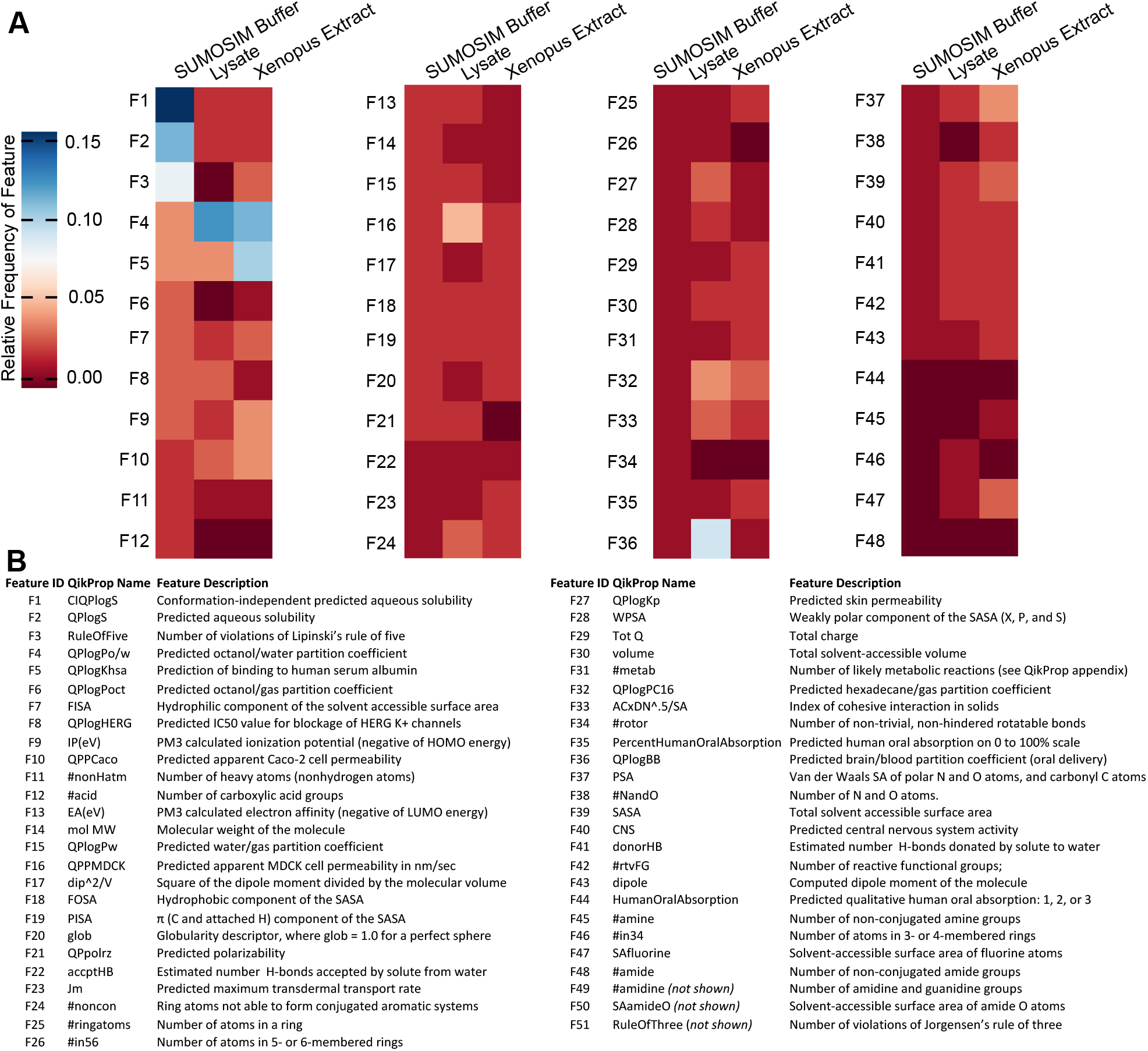
Heat map comparing molecular descriptor importance for SUMOSIM condensates in different solution environments. A. Relative feature importance in models of small molecules partitioning into SUMOSIM condensates in buffer, U2OS cell lysate, or concentrated *xenopus laevis* oocyte extract. B) Feature identifiers with the corresponding QikProp names and descriptions.

## List of tables

**Table S1**. Sequences of proteins and DNA molecules used in this study.

**Table S2**. Metabolites used in this study.

**Table S3**. Names and properties of FDA-approved drugs used in this study.

**Table S4**. Fluorescent drugs used in this study, and their spectroscopic properties.

**Table S5**. Partition coefficients of all compounds in all condensates used in this study.

**Table S6**. Physical properties of the scaffolds used to generate the four condensates examined in this study.

**Table S7**. Comparison of algorithms and feature sets used to generate models for this study. **Table S8**. XGBoost statistics for partitioning models of small molecules with the individual condensates used in this study.

**Table S9**. Parameters used to tune XGBoost models for this study. The XGBoost regression objective was set to minimize the squared error in predicted logPC. In addition to the tabulated tuning parameters, the following XGBoost parameters were consistent across all models in this study: minimum child weight = 4, maximum delta step = 2.

## Methods

### Lead Contact and Materials Availability

Requests for reagents should be directed to the Lead Contact, Michael K. Rosen. Plasmids used in this study are available by contacting the Lead Author.

### Genes, Plasmids, and DNA

polyPRM, polySH3, polySUMO, polySIM, Dhh1, GFP-Dhh1, and human cGAS were described previously ^6,21,22,33^. For detailed information, see supporting information. Sequences of proteins and DNA used in this study are listed in Table S1.

### Protein Expression, Purification, and Labeling

All proteins were expressed and purified from *E. coli* strain BL21 DE3T1R. Unless otherwise specified, all the purification steps were carried out at 4 °C. All proteins, except Dhh1, GFP-Dhh1, and cGAS, were purified using a similar protocol ^6,21^. For detailed information about each protein, see supporting information.

### Metabolites, drugs, and small molecule fluorophores

The metabolite library of 200 compounds (Table S2) used in this study covers most major metabolic pathways, including glycolysis, the tricarboxylic acid cycle, the pentose-phosphate pathway, and metabolism of amino acids and nucleotides. The drug library used is the Prestwick Chemical Library®:1520 FDA-approved & EMA-approved drugs (Prestwick chemical libraries (PCL1520.10-100-96G, Table S3).

### Measurement of protein PC values and droplet volume fraction by fluorescence microscopy

We measured protein partition coefficient (PC) values and calculated droplet volume fraction using confocal fluorescence microscopy with 1% Alexa-488 labeled polySUMO, polySH3, and cGAS, or 1% GFP-labeled Dhh1 in 25mM HEPES-NaOH (pH 7.4), 150 mM NaCl buffer.

### Microscopy assay to measure partitioning of fluorescent small molecules

To measure the partition coefficients of fluorescent small molecules using microscopy, we mixed unlabeled scaffold macromolecules (2-10 µM) with individual small molecules (100 nM-1 µM) in 25 mM HEPES-NaOH (pH 7.4), 150 mM NaCl buffer.

### Mass spectrometry assay to measure partitioning of small molecules

To measure the partition coefficients of fluorescent small molecules using mass spectrometry (MS) we mixed unlabeled macromolecular scaffolds with a final concentration of 2 µM of each metabolite (total of 200 metabolites, giving total metabolite concentration of 400 µM) or 1 µM of each drug (total of 300 drugs/sub-library, giving total drug concentration of 300 µM, with a final DMSO concentration of 3% v/v) in 25mM HEPES-NaOH (pH 7.4), 150 mM NaCl buffer. To generate SUMOSIM condensates in U2OS cell lysates, we modified a protocol used for biomimetic reconstitutions of stress granules and nucleoli^34^. Xenopus oocyte extracts were prepared using a standard protocol^35^ and flash frozen at -80 °C. Assays containing SUMOSIM condensates and drugs were generated and processed as for the U2OS cell lysates. We used a previously described protocol to quantify metabolites extracted from the different samples^36^. To detect and quantify drugs and other small molecules, we employed a standard untargeted metabolomic approach^37^. Mass spectrometric analyses were performed on Sciex QTRAP 6500+ mass spectrometer equipped with an electrospray ion (ESI) source (for targeted metabolomics) or a Sciex TripleTOF 6600 system (AB SCIEX, Framingham, MA, USA) equipped with electrospray ionization (ESI), atmospheric pressure chemical ionization (APCI) sources and calibrant delivery system (CDS) for the untargeted metabolomics approach).

### Isothermal Titration Calorimetry

ITC measurements were performed at 25 °C and 35 °C using a MicroCal PEAQ-ITC calorimeter (Malvern Panalytical). Purified polySUMO and polySIM were exchanged into 25 mM HEPES (pH 7.4) 150 mM NaCl buffer by size exclusion chromatography. Proteins were flash frozen in liquid nitrogen and stored at −80 °C. Immediately prior to measurement, proteins were thawed and diluted into the same buffer and mixed at a final concentration of 2 µM polySUMO and 2 µM polySIM (20 µM module concentrations), which is below the LLPS threshold. DMSO was also added to a concentration of 2% v/v, to match that of the drug solution in the syringe.

### Small Molecule Parameterization and Clustering

Low-energy 3D chemical structures of the small molecules were generated from SMILES strings (Tables S2 and S3) with LigPrep^38^ using the OPLS4 forcefield ^39^. The molecules were desalted, and protonation and tautomer states and were corrected for a pH 7.0 solution using Epik^40^. The structures were encoded as 51 ADME features using QikProp^23^. These small molecule parameters were used as the input to generate a UMAP chemical space map: an amenable UMAP embedding of the features was generated and the HSBCAN algorithm^24,25^ was applied to identify clusters. HDBSCAN assigned most small molecules to 10 defined clusters, and 6% of small molecules were not assigned to a cluster. Mordred descriptors were computed and curated for the relevant statistical models from SMILES strings of the small molecules using an in-house Python script. See supporting information for more information and to access relevant code.

### Statistical Modeling

Balanced raining, test, and validation sets for modeling were defined using the clustering analysis. Small molecules that were not assigned to a cluster were automatically assigned to the training set, then 20% of the molecules from each cluster were randomly selected for the validation set. Of the remaining molecules, 70% from each cluster were randomly selected as the training set and the final 30% were used as a test set. Small molecule descriptors were scaled and statistical models (and associated measured of performance) were generated using an in-house Python script. See supporting information for more information and to access relevant code.

## Acknowledgements

We thank Zhijian J Chen for providing the cGAS construct, Jeon Lee for initial help with modeling, Reshma Veettil for help with plotting heatmaps in Figure 1, William Peeples and Simon Curry for help with protein purification, and Rosen lab members for discussion. Research in the laboratory of M.K.R. is supported by the Howard Hughes Medical Institute, a Paul G. Allen Frontiers Distinguished Investigator Award and grants from the NIH (R35GM141736 and the Welch Foundation (I-1544). Research in the laboratory of M.S.S was supported by the NIH (R35 GM136271). This article is subject to HHMI’s Open Access to Publications policy. HHMI lab heads have previously granted a nonexclusive CC BY 4.0 license to the public and a sublicensable license to HHMI in their research articles. Pursuant to those licenses, the author-accepted manuscript of this article can be made freely available under a CC BY 4.0 license immediately upon publication.

## Author Contributions

M.K.R. and S.A.T. conceived of the study. S.A.T. performed biochemical experiments, H.D.C. performed computational analyses, H.B. performed mass spectrometry experiments, A.S.L. performed sub sample correlation analyses. S.A.T., H.D.C., A.S.L., M.S.S. and M.K.R. analyzed and interpreted the data. S.A.T., H.D.C., M.S.S. and M.K.R. drafted the manuscript, and all authors edited.

## Additional Information

Supplementary Information is available for this paper

Correspondence and requests for materials should be addressed to M.S.S. and M.K.R.

