## Supporting information for "Small Molecule Properties Define Partitioning into Biomolecular Condensates"

**Genes , Plasmids, and DNA**

polyPRM, polySH3, polySUMO, polySIM, Dhh1, GFP-Dhh1, and human cGAS were described previously [1-4]. All the genes, except human cGAS, were cloned into a modified pMAL plasmid (pMTTH) containing TEV-cleavable N-terminal MBP and C-terminal $His_6$ tags. Full-length human cGAS was cloned into a pSUMO bacterial expression vector containing Ulp1-cleavable N-terminal $His_6$ and SUMO tags. Sense and antisense single stranded DNAs (45bp) of a cGAS immunostimulatory DNA (ISD)[3] were purchased from integrated DNA technologies. Double-stranded ISD was generated by annealing the single stranded DNA oligonucleotides in 25 mM HEPES-NaOH pH 7.5, 50 mM NaCl ramping temperature from 95 °C to 25 °C at 1 °C/min [3]. Annealing efficiencies were > 99% as assessed by HPLC. Sequences of proteins and DNA used in this study are listed in Table S1.

**Protein Expression, Purification, and Labeling**

All proteins were expressed and purified from *E. coli* strain BL21 DE3T1R. Unless otherwise specified, all the purification steps were carried out at 4 °C. All proteins, except Dhh1, GFP-Dhh1, and cGAS, were purified using a similar protocol [1,2]. Transformed bacteria were grown to $OD_{600}$ of 0.6-0.8 and then induced with 1 mM IPTG at 18 °C for 16 hours. Cells were collected by centrifugation (4,700 x g, 30 minutes) and the pellet was resuspended in 50 mM Tris pH 8, 150 mM NaCl, 10 mM imidazole, and 5 mM β-mercaptoethanol (BME) with protease inhibitors. Cells were lysed using a cell disruptor (Emulsiflex-C5, Avestin), and lysates were cleared by centrifugation (45,000 x g, 45 minutes). Proteins were affinity-purified with Ni-NTA Agarose Resin (Qiagen), followed by Amylose Resin (NEB). The amylose eluate was digested with TEV protease (~1:100) overnight at 4 °C, filtered (0.22 μm) and loaded onto anion/cation exchange resin (Source15Q or Source 15S, GE Healthcare), then eluted with a linear gradient of NaCl (50–400 mM) in 50 mM Tris pH 8.0, 1 mM DTT and 1 mM EDTA. Protein-containing fractions were collected, concentrated, filtered, and further purified using size exclusion chromatography (Superdex 75 or 200, GE Healthcare) in 25 mM HEPES-NaOH pH 7.4, 150 mM NaCl, and 1 mM DTT buffer. After size exclusion chromatography, protein-containing fractions were concentrated by ultrafiltration (Amicon centricon) with 3k (polyPRM and polySIM) and 10k (polySH3 and polySUMO) molecular weight cutoffs. Single-use aliquots were flash frozen in liquid nitrogen and stored at -80 °C.

For Dhh1/GFP-Dhh1, the cell pellet was resuspended in lysis buffer (50 mM Tris pH 8, 500 mM NaCl, 10 mM imidazole, 5 mM BME with protease inhibitors). Cells were lysed using a cell disruptor (Emulsiflex-C5, Avestin), and lysates were cleared by centrifugation (45,000 x g, 45 minutes). Cleared lysate was incubated for 1 hour at 4 °C with Ni-NTA agarose resin (Bio-Rad) equilibrated with lysis buffer. Proteins were eluted by gravity chromatography using the following buffers. wash 1: 50 mM Tris pH 8, 500 mM NaCl, 10 mM imidazole, 5 mM BME; wash 2: 50 mM Tris pH 8, 100mM ATP, 2.5 M NaCl, 10 mM imidazole, and 5 mM BME; wash 3: 50 mM Tris pH 8, 10 mM imidazole, and 5 mM BME; elution: 50 mM Tris pH 8, 500 mM NaCl, 300 mM imidazole, and 5 mM BME. The eluate was added to amylose resin (New England Biolabs) and incubated for 30 minutes. Proteins were eluted by gravity chromatography using the following buffers. Amylose wash 1: 50 mM Tris pH 8, 150 mM NaCl, and 5 mM BME; amylose elution: 50

mM Tris pH 8, 150 mM NaCl, 5 mM BME, and 50 mM maltose. The amylose eluate was diluted threefold into 50 mM Tris pH 8, 5 mM BME buffer for a final concentration of 50 mM NaCl and filtered through a 0.22 μm Whatman filter (GE Healthcare). Filtrate was loaded onto a Source15Q ion-exchange column and eluted with a linear gradient of NaCl (50 – 500 mM) in 50 mM Tris pH 8.0, 1 mM DTT and 1 mM EDTA. Fractions containing the desired protein were collected, concentrated, filtered, and loaded onto a SD200 size exclusion column equilibrated with 25mM HEPES-NaOH pH 7.4, 150 mM NaCl, and 1 mM DTT buffer. After size exclusion chromatography, protein-containing fractions were concentrated by ultrafiltration (Amicon centricon) with 10k molecular weight cutoffs. Single-use aliquots were flash frozen in liquid nitrogen and stored at -80 °C.

For full-length human cGAS (hcGAS-FL) purification [3], the cell pellet was resuspended in lysis buffer (50 mM Tris pH 8, 500 mM NaCl, 10 mM imidazole, 5 mM BME with protease inhibitors). Cells were lysed using a cell disruptor (Emulsiflex-C5, Avestin), and lysates were cleared by centrifugation (45,000 x g, 45 minutes). Cleared lysate was incubated with Ni-NTA agarose resin (Bio-Rad) equilibrated with lysis buffer for 1 hour at 4 °C with circular rotation in 50 mL conical tubes. Proteins were eluted by gravity chromatography using the following buffers. wash 1: 50 mM Tris pH 8, 500 mM NaCl, 10 mM imidazole, 5 mM BME; wash 2: 50 mM Tris pH 8, 100mM ATP, 2.5 M NaCl, 10 mM imidazole, and 5 mM BME; wash 3: 50 mM Tris pH 8, 10 mM imidazole, and 5 mM BME; elution: 50 mM Tris pH 8, 300 mM NaCl, 300 mM imidazole, and 5 mM BME. The eluate was digested with Ulp (~1:100) overnight at 4 °C, filtered through a 0.22 μm Whatman filter (GE Healthcare) and loaded onto a HiTrap Heparin column (GE Healthcare), then eluted with a linear gradient of NaCl (50–500 mM) in 50 mM Tris pH 8.0, 1 mM DTT and 1 mM EDTA. Protein-containing fractions were collected, concentrated, filtered, and further purified using size exclusion chromatography using an SD200 size exclusion column (GE Healthcare) ) in 25 mM HEPES-NaOH, pH 7.4, and 150 mM NaCl. After size exclusion chromatography, protein-containing fractions were concentrated by ultrafiltration (Amicon centricon) with 10k molecular weight cutoffs. Single-use aliquots were flash frozen in liquid nitrogen and stored at -80 °C.

For all proteins, purity was assessed at each step of purification using SDS-PAGE. For experiments requiring protein fluorescence (protein PC value and droplet volume measurements using confocal fluorescence microscopy), all proteins except Dhh1 (we used GFP-Dhh1 for protein fluorescence) were labeled using maleimide-conjugated Alexa 488 dye (Life Technologies) following the manufacturer's protocol. After labeling, proteins were separated from free dye on an SD200 size exclusion column (GE Healthcare) in 25 mM HEPES-NaOH, pH 7.4, and 150 mM NaCl, and concentrated by ultrafiltration (Amicon centricon, 10k molecular weight cutoff). As assessed by UV-Vis spectrophotometry, all proteins were labeled at approximately 95%. Single-use aliquots were flash frozen in liquid nitrogen and stored at -80 °C.

**Metabolites, drugs, and small molecule fluorophores**

The metabolite library of 200 compounds (Table S2) used in this study is a calibration standard for targeted metabolomics, which covers most major metabolic pathways, including glycolysis, the tricarboxylic acid cycle, the pentose-phosphate pathway, and metabolism of amino acids and nucleotides. A metabolite stock (50 µM of each compound) was prepared in MS grade water (Sigma-Aldrich) and used in partitioning experiments to give a final concentration of 2 µM for each compound. The drug library used is the Prestwick Chemical Library®:1520 FDA-approved & EMA-approved drugs (Prestwick chemical libraries (PCL1520.10-100-96G, Table S3). We purchased drugs as individual 10 mM stock solutions in DMSO. We mixed compounds to prepare sub-libraries of ~300 molecules of unique molecular weight (30 μM of each drug in MS grade water/DMSO 10/90% by volume. For the drug partitioning experiments, the sub-libraries were diluted 30-fold into the reaction mixture, producing a final concentration of ~1 μM of each compound and ~3% DMSO. Using a ChemiDoc XRS+ system (Bio-Rad) at different excitation wavelengths, 488 nm, 546 nm, and 647 nm, we identified 34 fluorescent molecules in the drug library (Table S4). We prepared individual working stock solutions of these molecules (200 μM in MS grade water/DMSO, 98/2 % by volume) for use in partitioning experiments using confocal fluorescence microscopy. To optimize our methods to quantify small molecule partitioning into biomolecular condensates, including extraction of compounds from droplet and bulk samples (see below), we used commonly available fluorophores, such as FITC, Rhodamine, Alexa488, etc. (Table S4). We prepared individual working stock solutions of these fluorophores (100 μM in MS grade water/DMSO, 99/1 % by volume) for use in partitioning experiments using confocal fluorescence microscopy and MS.

**Microscopy plate preparation**

Microscopy experiments were carried out in 384-well glass bottom microwell plates (Brooks Life Science Systems: MGB101-1-2-LG-L). Prior to use, plates were washed with 5% Hellmanex at 37°C for 4 h and then extensively with MilliQ water. Glass was etched with 1 M NaOH for 1 h at room temperature, washed extensively with MilliQ water, and then treated overnight (≥16 h) at room temperature with 25 mg/mL 5K mPEG-silane (PEGWorks) in 95% Ethanol. The plate was washed once with 95% ethanol, extensively with MilliQ water, and then dried in a chemical hood for 3-4 h. PEGylated microscopy plates were sealed with adhesive PCR plate foil (Thermo). Immediately prior to use, foil was cut above individual wells and both plastic and PEGylated glass were passivated by incubation with freshly prepared 10 mg/mL BSA for 30 min. Wells were rinsed once with MilliQ water, followed by buffer (25mM HEPES-NaOH (pH 7.4) with 150 mM NaCl) to remove excess BSA, and microscopy samples (50-60 μL) were immediately added. Desiccation of microscopy samples was minimized following transfer to the plate by sealing with transparent sealant tape.

**Measurement of protein PC values and droplet volume fraction by fluorescence microscopy**

We measured protein partition coefficient (PC) values and calculated droplet volume fraction using confocal fluorescence microscopy with 1% Alexa-488 labeled polySUMO, polySH3, and cGAS, or 1% GFP-labeled Dhh1 in 25mM HEPES-NaOH (pH 7.4), 150 mM NaCl buffer. We used the following concentrations of each species: 5 μM polySUMO (module concentration of 50 μM) and 5 μM polySIM (module concentration of 50 μM); 10 μM polySH3 (module concentration of 50 μM) and 10 μM (polyPRM (module concentration of 50 μM); 5 μM MBP-Dhh1 and 0.2 μM TEV-protease; 2 μM cGAS and 2 μM 45bp DNA. The scaffold mixtures were incubated for 1 hour (cGAS-DNA), 4 hours (SUMO/SIM), 12 hours (SH3/PRM), or 20 hours (Dhh1) at room temperature. After incubation, we acquired images using a 20x air objective (for SUMO/SIM and SH3/PRM) or 60x oil immersion objective (for Dhh1 and cGAS-DNA) on a Leica SP8 Laser Scanning Confocal Microscope.

To avoid dilution effects of the microscope point spread function (PSF) on intensities of smaller condensates, for partition coefficient and droplet volume measurements, we only analyzed droplets with x-y diameter >2-fold larger than the z-dimension PSF. Intensities from all included regions of droplets or bulk phases were separately averaged and used to calculate concentrations in the two phases using intensity vs. concentration standard curves. Standard curve solutions were prepared alongside experimental samples using GFP-Dhh1 or Alexa 488-labeled molecules (polySUMO, polySH3, and cGAS) in the corresponding experimental buffer (25mM HEPES-

NaOH (pH 7.4), 150 mM NaCl) supplemented with 0.1-0.2 mg/mL BSA to prevent fluorophore adsorption to surfaces. Partition coefficients were determined from the bulk and droplet concentrations.

$$\text{Partition coefficient} = \frac{Droplet\ concentration}{Bulk\ concentration}$$

Droplet volume fraction was determined from the total, droplet and bulk concentrations by [5]:

$$\text{Droplet volume fraction} = \frac{(Total\ concentration - Bulk\ concentration)}{(Droplet\ concentration - Bulk\ concentration)}$$

**Microscopy assay to measure partitioning of fluorescent small molecules**

To measure the partition coefficients of fluorescent small molecules using microscopy, we used the following concentrations of unlabeled scaffold macromolecules mixed with individual small molecules (100 nM-1 μM) in 25 mM HEPES-NaOH (pH 7.4), 150 mM NaCl buffer: 5 μM polySUMO (module concentration of 50 μM) and 5 μM polySIM (module concentration of 50 μM); 10 μM polySH3 (module concentration of 50 μM) and 10 μM polyPRM (module concentration of 50 μM); 5 μM MBP-Dhh1 and 0.2 μM TEV-protease; 2 μM cGAS and 2 μM 45bp DNA. The macromolecules were added to prepared 384-well plates first, followed by the fluorescent small molecules; mixtures were incubated for 1 hour (cGAS-DNA), 4 hours (SUMO/SIM), 12 hours (SH3/PRM), or 20 hours (Dhh1) at room temperature. After incubation, we acquired images using a 20x air objective (for SUMO/SIM and SH3/PRM) or 60x oil immersion objective (for Dhh1 and cGAS-DNA) on a Leica SP8 Laser Scanning Confocal Microscope. Fluorescence intensity (*I*) from the droplet and bulk phases were used to measure the partition coefficients. As a background control, an equal volume of DMSO was added to the unlabeled scaffold mixture, and measured values in the droplet and solution phases were used in calculation of partition coefficients according to:

$$\text{Partition coefficient} = \frac{(I_{Droplet(drug)} - I_{Droplet(DMSO)})}{(I_{bulk(drug)} - I_{bulk(DMSO)})}$$

**Mass spectrometry assay to measure partitioning of small molecules**

***Extraction of small molecules from droplet and bulk samples***

The mass spectrometry (MS) assay required efficient extraction of small molecules from condensates and bulk solutions. To optimize this process[6], we first measured partitioning of small molecule fluorophores (Table S4) into the SH3PRM condensate using confocal fluorescence microscopy as a gold standard, as described above. We then compared this value to those measured by fluorescence spectroscopy and MS, which both required extraction of the dye from droplet and bulk solutions. To produce these samples, we incubated SH3PRM condensates with FITC, and then separated droplets from bulk by centrifugation as detailed in the following section. To extract the dye from the samples we first digested the protein scaffolds with trypsin, varying time (1-12 hours), temperature (25 °C or 37 °C), and enzyme concentration (0.01-5 µg/µL). We used methanol (80:20 v/v solvent:sample) to quench the reaction and precipitate the digested proteins at different temperatures (4 °C, -20 °C, -80 °C) and for different durations (4-24 hours). For each combination of treatments, we quantified the amount of dye present in the droplet and bulk fractions by fluorescence spectroscopy ($\lambda_{ex}$ = 488 nm, $\lambda_{em}$ = 495-600 nm) through a standard curve. In parallel, we also measured the amount of extracted FITC by MS as detailed below. Processing conditions were optimized to reproducibly yield identical PC values by all three methods. Final conditions are described in the sections below. We note that tryptic digestion prior to methanol precipitation was essential to achieving complete extraction of dye from the condensate samples.

***Sample preparation for condensates reconstituted in simple buffers***

For small molecule partitioning experiments using mass spectrometry (MS) we mixed macromolecular scaffolds with a final concentration of 2 μM of each metabolite (total of 200 metabolites, giving total metabolite concentration of 400 μM) or 1 μM of each drug molecule (total of 300 drug molecules/sub-library, giving total drug concentration of 300 μM, with a final DMSO concentration of 3% v/v) in 25mM HEPES-NaOH (pH 7.4), 150 mM NaCl buffer. Scaffolds were used at the following concentration: 5 μM polySUMO (module concentration of 50 μM) and 5 μM polySIM (module concentration of 50 μM); 10 μM polySH3 (module concentration of 50 μM) and 10 μM polyPRM (module concentration of 50 μM); 5 μM MBP-Dhh1 and 0.2 μM TEV-protease; 2 μM cGAS and 2 μM 45bp DNA. In each case macromolecules were mixed first, followed by small molecules. Mixtures were incubated for 1 hour (cGASDNA), 4 hours (SUMOSIM), 12 hours (SH3PRM), or 20 hours (Dhh1) at room temperature. Total volume of each reaction was 1000 μL. For were replicated twice and 8-9 times for drugs and metabolites, respectively. We also prepared the following control samples for each condensate: (1) macromolecules without any added small molecules, (2) small molecules without any macromolecules, and (3) single macromolecule component of condensates with small molecules (e.g. polySUMO or polySIM alone with small molecules; in all such samples phase separation did not occur). After incubation, samples were centrifuged at 14,000 g for 30 min using a temperature-controlled centrifuge maintained at 22 °C.

The supernatant was transferred carefully and as completely as possible to a new 1.5 mL centrifuge tube, leaving the condensate pellet (typically ~10 μL, for 1000 μL total reaction volume) in the original tube. Based on the droplet volume measurements (see above), we transferred an equal droplet volume of the supernatant into a new micro centrifuge tube. The supernatant and pellet fractions were each mixed with trypsin (10 μL, from 0.1 μg/μL stock concentration MS-grade, Porcine; Fisher Scientific) prepared in 25mM HEPES-NaOH (pH 7.4), 150 mM NaCl buffer in MS grade water, and incubated at 37 °C for 12 h with gentle shaking. After incubation, the reactions were mixed well by pipetting, and 80 μL of MS grade methanol pre-chilled to -80 °C was added (to give 80% v/v methanol). Samples were vortexed for 30 s and incubated at -80 °C for 12 hours. After incubation, samples were vortexed again for 30 s and centrifuged at 20,000 g at 22 °C for 45 minutes. After centrifugation, supernatant fractions containing small molecules were carefully removed and kept at -80°C prior to MS analysis.

***Sample preparation for condensates reconstituted in U2OS cell lysates, and Xenopus extracts***

To generate SUMOSIM condensates in U2OS cell lysates, we modified a protocol used for biomimetic reconstitutions of stress granules and nucleoli [7]. U2OS cells were cultured in DMEM ((HyClone) supplemented with 10% FBS (HyClone; SH30071.03 and SH30396.03) and maintained at 37°C in a humidified incubator with 5% $CO_2$. Cells were grown to 100% confluency in 10-cm cell culture–treated dishes, the medium was aspirated, and cells were washed with PBS. Cells were detached by scraping in 5 ml PBS and recovered by centrifugation at 500 *g* for 3-5 min. Buffer was carefully removed by aspiration and cell pellets were stored at −80°C. To prepare lysate, pellets (10-15 μL volume) were resuspended in 250 μL cell lysis buffer (25 mM HEPES-NaOH (pH 7.4), 100 mM NaCl with protease inhibitors (Roche 11836170001)), and pipetted 5-10 times using a 200 μL tip to produce a visibly homogenous solution, followed by incubation on ice for 10 min. The suspension was freeze-thawed five times by alternating between liquid nitrogen (15 sec) and a 37° water bath (2 min), and finally pipetted twice using a 200 μL tip. The lysate was cleared by centrifugation at 20,000 g for 5 min at 4 °C. The supernatant was transferred to a fresh tube and used within 1-2 hours. Total protein concentration in the cleared lysate was estimated as 4 mg/ml using the Bradford assay (Bio-Rad, Cat# 5000002). To generate condensates in this system, unlabeled polySUMO and polySIM were added to cleared cell lysate to final concentrations of 10 μM, followed by 1 μM each drug molecule (10 μL of 30 μM stock of 300 drug molecules/sub-library), giving a final assay volume of 280 μL (final composition of 3% v/v DMSO). We also prepared control samples containing drug molecules in lysis buffer or cell lysate alone (without polySUMO or polySIM proteins). After drug addition, mixtures were incubated at room temperature for 4 hours, and droplet and solution phases were separated by centrifugation at 2000 g for 15 min at 22°C. Note that these conditions were optimized to minimize background sedimentation of drugs with the lysate only control, while still recovering most of condensates. The supernatant was transferred carefully and as completely as possible to a new 1.5 mL centrifuge tube, leaving the condensate pellet (typically ~5 μL) in the original tube. Based on the droplet volume measurements (see above), we transferred an equal droplet volume of the supernatant into a new centrifuge tube. The supernatant and droplet fractions were processed and prepared for MS as described above for condensates in simple buffers.

Xenopus oocyte extracts were prepared using a standard protocol[8] and flash frozen at -80 °C. To minimize background sedimentation of drugs in the absence of condensates, we clarified the raw extract by ultracentrifugation at 100,000 g for 1 hour at 4 °C prior to use. Total protein concentration in the cleared extract was estimated as ~80 mg/ml using the Bradford assay (Bio-Rad, Cat# 5000002). Assays containing SUMOSIM condensates and drugs were generated and processed as for the U2OS cell lysates.

***Targeted Metabolomics approach to quantify the partitioning of metabolites***

We used a previously described protocol to quantify metabolites extracted from the different samples[9]. LC-MS/MS mass spectrometric analyses were performed on a Sciex QTRAP 6500+ mass spectrometer equipped with an electrospray ion (ESI) source. The ESI source was used in both positive and negative ion modes, configured as follows: Ion Source Gas 1 (Gas 1), 40psi; Ion Source Gas 2 (Gas 2), 35 psi; curtain gas (CUR), 50 psi in the negative polarity mode and 45 psi in the positive polarity mode; source temperature, 550 °C; and ion spray voltage (IS), +4800 V(+) and −4000 V (−). The mass spectrometer was coupled to a Shimadzu HPLC (Nexera X2 LC-30AD). The system is controlled by Analyst 1.7.2 software.

Hydrophilic interaction chromatography was performed using a SeQuant® ZIC®-pHILIC 5 μm polymeric 150 × 2.1 mm PEEK coated HPLC column (Millipore Sigma, USA). The column temperature, sample injection volume, and flow rate were 45 °C, 5 μL, and 0.15 mL/min respectively. HPLC conditions were as follows: Solvent A: 20 mM ammonium carbonate including 0.1% Ammonium hydroxide. Solvent B: Acetonitrile. Gradient pattern was 0 min: 80% B, 20 min: 20% B, 20.5 min 80% B, 34 min: 80% B. The mass spectrometer was equipped with a switching valve such that the column effluent between 0-2 min and 18-34 min was delivered to the waste to avoid interfering ions entering the ESI source in the negative polarity mode. In the positive polarity mode, the column effluent between 0-1min and 18-34 min was delivered to waste. All targeted metabolites were eluted from the column between 2-15 minutes. Data were processed using SCIEX OS 2.1 software with relative quantification based on the peak area of each metabolite.

***Untargeted metabolomic approach to quantify partitioning of drugs and other small molecules***

To detect and quantify drugs and other small molecules such as fluorophores, we employed a standard untargeted metabolomic approach[10]. Mass spectrometric analyses were performed on a

Sciex TripleTOF 6600 system (AB SCIEX, Framingham, MA, USA) equipped with electrospray ionization (ESI) , atmospheric pressure chemical ionization (APCI) sources and calibrant delivery system (CDS). The electrospray ionization (ESI) source used in the positive and negative ionization mode and configured as follows: Ion Source Gas 1 (Gas 1), 50psi; Ion Source Gas 2 (Gas 2), 45 psi; curtain gas flow, 35 psi; source temperature, 550 °C; and ion spray voltage floating, +5500 V(+) and −4500 V (−). TOF-MS mode (Full scan) and Information Dependent Acquisition (IDA) mode (Product Ion scan) were utilized to collect MS and MS/MS data, respectively. For TOF-MS scans, the mass range was from m/z 70 to 1000 and for Product Ion scans, the mass range was from m/z 30 to 1000. The collision energy (CE) was set at 30 V (+) or -30 V (-) and collision energy spread (CES) was ±15 V. The accumulation time was 0.25 seconds for TOF-MS scans and 0.05 seconds for product ion scans. The instrument was automatically calibrated for mass accuracy (<5 ppm), including MS1 scan and MS/MS scan, every five samples using APCI calibration solution. The mass spectrometer was coupled to a Shimadzu HPLC (Nexera X2 LC-30AD). The system was controlled by Analyst TF 1.8.1 software (Sciex).

Reverse phase chromatography was performed using an ACE 3 C18-PFP 150 x 4.6 mm HPLC column (Mac-Mod, USA). The column temperature, sample injection volume, and flow rate were 30°C, 10 µL, and 0.5 mL/min respectively. The HPLC conditions were as follows: Solvent A: Water with 0.1% Formic Acid (v/v), LC/MS grade and Solvent B: Acetonitrile with 0.1% Formic Acid (v/v), LC/MS grade. Gradient condition was 0-2 min: 5% B, 5-16 min: 90% B, 17 min 5% B, 30 min: 5% B. Compounds eluted in a linear gradient of 5% to 90% B over 14 minutes were used for the analysis. Data were processed using SCIEX OS 2.1 software with relative quantification based on the peak area of each small molecule.

For samples prepared in simple buffers, we did not observe any measurable amounts of metabolites or small molecules in the pellet fraction of control samples (without condensates). Hence, we determined Partition coefficient (PC) values simply by dividing the integrated peak areas for each molecule in the droplet and bulk samples:

$$PC = \frac{Peak\ area_{Droplet}}{Peak\ area_{bulk}}$$

In the U2OS lysate and Xenopus extract solutions used with the SUMOSIM system, many of the compounds did sediment substantially in control samples lacking condensates. To minimize the errors generated by this background, we discarded compounds where the amount of material sedimented in the lysate/extract alone was > 50% of that sedimented in the presence of SUMOSIM condensates. For all other compounds, the background sedimentation was corrected in calculating PC according to:

$$PC = \frac{Peak\ area_{pellet}(SUMOSIM + lysate) - \ Peak\ area_{pellet}(lysate\ alone)}{Peak\ area_{supernatant}(SUMOSIM + lysate)}$$

**Isothermal Titration Calorimetry**

ITC measurements were performed at 25 °C and 35 °C using a MicroCal PEAQ-ITC calorimeter (Malvern Panalytical). Purified polySUMO and polySIM were buffer exchanged by size exclusion chromatography on a Superdex200 column with a mobile phase buffer of 25 mM HEPES buffer (pH 7.4) and 150 mM NaCl. Protein concentrations were measured using UV-Vis spectrophotometry with calculated extinction coefficients, $\varepsilon_{280nm,polySUMO}$ = 27,390 $M^{-1}cm^{-1}$ and $\varepsilon_{280nm,polySIM}$ = 6,990 $M^{-1}cm^{-1}$. Proteins were flash frozen in liquid nitrogen and stored at −80 °C. Immediately prior to measurement, proteins were thawed and diluted into the same buffer and mixed at a final concentration of 2 µM polySUMO and 2 µM polySIM (20 µM module concentrations) , which is below the LLPS threshold. DMSO was also added to a concentration of 2% v/v, to match that of the drug solution in the syringe. For each data point, 1.9 µL of 200 µM drug in the same buffer were injected into 0.3 ml of protein every 120 s. Twenty injections were performed per experiment. We also acquired analogous control thermograms where drug molecules were injected into buffer, or where buffer was injected into protein. The latter did not produce significant heat changes and was not used for further correction of data. However, injection of some drug molecules into buffer produced significant heat changes. In these cases, the control thermogram was subtracted from the drug + protein thermograms prior to analysis. Data for raw ITC and thermodynamic curves were analyzed using Microcal PEAQ-ITC software and plotted using SigmaPlot software.

**PRODAN-dye based condensate polarity measurements**

We used the solvatochromatic dye, 1-[6-(Dimethylamino)naphthalen-2-yl]propan-1-one (PRODAN), to characterize the polarity of the four biomolecular condensates studied here. We used confocal fluorescence microscopy (Leica SP8, $\lambda_{ex}$ 405 nm) to collect 5 nm wide emission spectral window measurements between 420 nm to 650 nm for the droplet and bulk phases. The wavelength of maximum emission intensity was estimated visually and could be compared to those obtained for PRODAN dissolved in pure solvents of different polarities, such as acetonitrile, DMSO, ethanol, methanol, and water.

**Sub sample correlation analysis**

As a qualitative depiction of the range of PC values at which correlations in PCs between the different condensate systems emerges, we developed an algorithm to characterize how correlations vary across different subsamples of the PC data. We subsample the data in multiple overlapping windows with a normally distributed bias, while varying the mean and standard deviation of the biasing normal distributions. We calculate Pearson correlation coefficients for the data in each window, then display the correlation coefficients as a raster image where x-axis position represents the center of the sampling window (i.e., the mean of the normal distribution that biases the sampling), and the y-axis position represents the size of the sampling window (i.e., the standard deviation). Note that if only two PC value pairs are sampled, the correlation coefficient will be 1 regardless of data values and is thus uninformative and we do not show them on our plots. The algorithm in pseudocode is below and R code[11] implementing the algorithm is provided as Supplementary Software.

**input: PC data vector for condensate 1** $D_1$ (length $n$),

PC data vector for condensate 2 $D_2$ (length $n$),

$j$ center positions of sampling windows $m_1, \ldots, m_j$

$k$ sampling window sizes (standard deviations) $s_1, \ldots, s_k$,

$i$ sampling iterations 1, …, $i$

convergence tolerance $t$

**output:** **5 ×** ($j$×$k$) matrix where the rows contain the 0, 0.25, 0.50, 0.75, and 1 quantiles of the sampled correlation coefficients and the columns correspond to each combination of $m$ and $s$

create empty vector $A$ # *for quantiles of B*

**for each** $m$:

  **for each** $s$:

    create empty vector $B$ # *for correlation coefficients*

    **for each** $i$:

      create empty vector $C$ # *for samples*

      create emtpy vector $D$ # *for fraction sampled*

      **while** tolerance not met:

obtain sample $d$ from ($D_1$, $D_2$)

obtain sample $x$ from Normal(0, $s$) distribution

**if** $|d_1-m| \leq |x|$ and $d$ was not previously sampled:

append $d$ to $C$

append length($B$)/$n$ to $D$

**if** last $t$ elements of $B$ are equal to the final element of $B$:

end while loop

append correlation coefficient of samples in C to B

append vector ($q_0$, $q_{0.25}$, $q_{0.5}$, $q_{0.75}$, $q_1$) to $A$ (where the $q$s are quantiles of B)

We set $j$ and $k$ to 100, $i$ to 100, and $t$ to 100. Running in parallel on 6 48-core server nodes the calculations take approximately 6 h.

**Small Molecule Parameterization**

Low-energy 3D chemical structures of the small molecules used in this study (N = 1507) were generated from SMILES strings (Tables S2 and S3) with LigPrep[12] using the OPLS4 forcefield[13]. The molecules were desalted, and protonation and tautomer states and were corrected for a pH 7.0 solution using Epik[14]. The structures were encoded as 51 ADME features using QikProp[15]; input files containing the QikProp descriptors used to build the UMAP chemical space and statistical models are available on the Sigman Lab Github: https://github.com/SigmanGroup/small-molecule-partitioning. Mordred descriptors were computed and curated for the relevant statistical models from SMILES strings of the small molecules using an implementation of the Mordred package which is available on the Sigman Lab Github: https://github.com/SigmanGroup/Mordred-descriptors.

**Chemical Space Analysis and Clustering**

The QikProp descriptors were used as the input to generate a 2D UMAP chemical space[16]: an amenable UMAP embedding of the features was generated and the HSBCAN algorithm was applied to identify clusters[17]. HDBSCAN is an unsupervised clustering method that automatically identified and assigned molecules to 10 clusters. HDBSCAN also identified that 6% of small molecules were not clustered because they did not overlap in UMAP space with defined clusters. Code to reproduce the UMAP chemical space and associated clustering (including the specific algorithm parameters used in study) is available on the Sigman Lab Github: https://github.com/SigmanGroup/small-molecule-partitioning.

**Statistical Modeling**

Training, test, and validation sets for modeling were designed as described in the Methods section of the main text. The descriptors were scaled by removing the mean and scaling to unit variance. Using training set statistics ($R^2$ and mean absolute error, MAE) as a benchmark, we assessed several tree-based statistical models, including random forest (RF) regressors, RF regressors with cost complexity pruning (CCP), and XGBoost models. Additionally, we compared the tree-based methods to a multilayer perceptron neural net. The neural net preformed worse than the tree-based methods, with the XGBoost model achieving the overall best performance (Table S7). For our initial survey of modeling algorithms, default parameters were used except as indicated in the attached code. RF and RF with CCP models were developed from ensembles of 1000 trees; for the models with CCP, the complexity parameter (cpp_alpha) was set to 0.001. Code to reproduce the statistical models in this study (including feature scaling and feature importance metrics) is available on the Sigman Lab Github: https://github.com/SigmanGroup/small-molecule-partitioning.

We also determined that the use of QikProp ADME-type descriptors resulted in better models than Mordred descriptors, which are more structurally based. Combining the two descriptor sets did not improve models, even if collinear descriptors ($R^2 > 0.9$) were eliminated or if descriptors were selecting selected using a *Z*-test mean comparison for excluded vs. included molecules using the statsmodels Python package (Table S7)[18]. Validation set molecules were not used in *Z*-test comparisons.

The XGBoost algorithm with QikProp features was implemented to make four separate but similar models of small molecules partitioning with SUMOSIM, SH3PRM, Dhh1, of cGASDNA condensates (Table S8, entries **1-4**). As described in the main text, an all condensate and an average LogPC model were also developed (Table S8, entries **5** and **6**). For the all condensate model, the experimental condensate Prodan measurements were added to the small molecule feature set. The XGBoost algorithm parameters were tuned separately for each of the six models as described in Table S9. XGBoost parameters for the models of small molecule partitioning into SUMOSIM condensates in U2OS cell lysate and Xenopus *laevis* oocyte extract are also indicated in Table S9. For validation of the average logPC model, the training and test sets were pooled (N pooled = 1145); the XGBoost model was retrained on the pooled without re-tuning of algorithm parameters and used to predict the average logPC for validation set molecules (N validation = 263).
