## Supplemental Table 1 for "Small Molecule Properties Define Partitioning into Biomolecular Condensates"

**Table S1: List of constructs, DNA molecules used in this study.**

| **No:** | **Name** |
| --- | --- |
| **1** | **polySUMO (SUMO-10R)**  **MGGSWGGSMSEEKPKEGVKTENDHINLKVAGQDGSVVQFKIKRHTPLSKLMKAYSERQGLSMRQIRFRFDGQPINETDTPAQLEMEDEDTIDVFQQQTVVGGSGGSGGSGGSMSEEKPKEGVKTENDHINLKVAGQDGSVVQFKIKRHTPLSKLMKAYSERQGLSMRQIRFRFDGQPINETDTPAQLEMEDEDTIDVFQQQTVVGGSGGSGGSGGSMSEEKPKEGVKTENDHINLKVAGQDGSVVQFKIKRHTPLSKLMKAYSERQGLSMRQIRFRFDGQPINETDTPAQLEMEDEDTIDVFQQQTVVGGSGGSGGSGGSMSEEKPKEGVKTENDHINLKVAGQDGSVVQFKIKRHTPLSKLMKAYSERQGLSMRQIRFRFDGQPINETDTPAQLEMEDEDTIDVFQQQTVVGGSGGSGGSGGSMSEEKPKEGVKTENDHINLKVAGQDGSVVQFKIKRHTPLSKLMKAYSERQGLSMRQIRFRFDGQPINETDTPAQLEMEDEDTIDVFQQQTVVGGSGGSGGSWGGSMSEEKPKEGVKTENDHINLKVAGQDGSVVQFKIKRHTPLSKLMKAYSERQGLSMRQIRFRFDGQPINETDTPAQLEMEDEDTIDVFQQQTVVGGSGGSGGSGGSMSEEKPKEGVKTENDHINLKVAGQDGSVVQFKIKRHTPLSKLMKAYSERQGLSMRQIRFRFDGQPINETDTPAQLEMEDEDTIDVFQQQTVVGGSGGSGGSGGSMSEEKPKEGVKTENDHINLKVAGQDGSVVQFKIKRHTPLSKLMKAYSERQGLSMRQIRFRFDGQPINETDTPAQLEMEDEDTIDVFQQQTVVGGSGGSGGSGGSMSEEKPKEGVKTENDHINLKVAGQDGSVVQFKIKRHTPLSKLMKAYSERQGLSMRQIRFRFDGQPINETDTPAQLEMEDEDTIDVFQQQTVVGGSGGSGGSGGSMSEEKPKEGVKTENDHINLKVAGQDGSVVQFKIKRHTPLSKLMKAYSERQGLSMRQIRFRFDGQPINETDTPAQLEMEDEDTIDVFQQQTVVGGSGGSENLYFQ** |
| **2** | **polySIM (SIM-10R)**  **MGGSWGGSKVDVIDLTIESSSDEEEDPPAKRGGSGGSGGSGGSKVDVIDLTIESSSDEEEDPPAKRGGSGGSGGSGGSKVDVIDLTIESSSDEEEDPPAKRGGSGGSGGSGGSKVDVIDLTIESSSDEEEDPPAKRGGSGGSGGSGGSKVDVIDLTIESSSDEEEDPPAKRGGSGGSGGSGGSKVDVIDLTIESSSDEEEDPPAKRGSKVDVIDLTIESSSDEEEDPPAKRGGSGGSGGSGGSKVDVIDLTIESSSDEEEDPPAKRGGSGGSGGSGGSKVDVIDLTIESSSDEEEDPPAKRGGSGGSGGSGGSKVDVIDLTIESSSDEEEDPPAKR** |
| **3** | **PolySH3 (SH3-5R)**  **MDLNMPAYVKFNYMAEREDELSLIKGTKVIVMEKSSDGWWRGSYNGQVGWFPSNYVTEEGDSPLASGAGGSEGGGSEGGTSGATDLNMPAYVKFNYMAEREDELSLIKGTKVIVMEKSSDGWWRGSYNGQVGWFPSNYVTEEGDSPLASGAGGSEGGGSEGGTSGATHMDLNMPAYVKFNYMAEREDELSLIKGTKVIVMEKSSDGWWRGSYNGQVGWFPSNYVTEEGDSPLASGAGGSEGGGSEGGTSGATDLNMPAYVKFNYMAEREDELSLIKGTKVIVMEKSSDGWWRGSYNGQVGWFPSNYVTEEGDSPLASGAGGSEGGGSEGGTSGATDLNMPAYVKFNYMAEREDELSLIKGTKVIVMEKSSDGWWRGSYNGQVGWFPSNYVTEEGDSPLGGGSENLYFQ** |
| **4** | **polyPRM (PRM-5R)**  **KGGSWGGSKKKKTAPTPPKRSGGSGGSGGSGGSKKKKTAPTPPKRSGGSGGSGGSGGSKKKKTAPTPPKRSGGSGGSGGSGGSKKKKTAPTPPKRSGGSGGSGGSGGSKKKKTAPTPPKRSGGSGSENLYFQ** |
| **5** | **Dhh1**  **MGSINNNFNTNNNSNTDLDRDWKTALNIPKKDTRPQTDDVLNTKGNTFEDFYLKRELLMGIFEAGFEKPSPIQEEAIPVAITGRDILARAKNGTGKTAAFVIPTLEKVKPKLNKIQALIMVPTRELALQTSQVVRTLGKHCGISCMVTTGGTNLRDDILRLNETVHILVGTPGRVLDLASRKVADLSDCSLFIMDEADKMLSRDFKTIIEQILSFLPPTHQSLLFSATFPLTVKEFMVKHLHKPYEINLMEELTLKGITQYYAFVEERQKLHCLNTLFSKLQINQAIIFCNSTNRVELLAKKITDLGYSCYYSHARMKQQERNKVFHEFRQGKVRTLVCSDLLTRGIDIQAVNVVINFDFPKTAETYLHRIGRSGRFGHLGLAINLINWNDRFNLYKIEQELGTEIAAIPATIDKSLYVAENDETVPVPFPIEQQSYHQQAIPQQQLPSQQQFAIPPQQHHPQFMVPPSHQQQQAYPPPQMPSQQGYPPQQEHFMAMPPGQSQPQY** |
| **6** | **Human cGAS**  **MQPWHGKAMQRASEAGATAPKASARNARGAPMDPTESPAAPEAALPKAGKFGPARKSGSRQKKSAPDTQERPPVRATGARAKKAPQRAQDTQPSDATSAPGAEGLEPPAAREPALSRAGSCRQRGARCSTKPRPPPGPWDVPSPGLPVSAPILVRRDAAPGASKLRAVLEKLKLSRDDISTAAGMVKGVVDHLLLRLKCDSAFRGVGLLNTGSYYEHVKISAPNEFDVMFKLEVPRIQLEEYSNTRAYYFVKFKRNPKENPLSQFLEGEILSASKMLSKFRKIIKEEINDIKDTDVIMKRKRGGSPAVTLLISEKISVDITLALESKSSWPASTQEGLRIQNWLSAKVRKQLRLKPFYLVPKHAKEGNGFQEETWRLSFSHIEKEILNNHGKSKTCCENKEEKCCRKDCLKLMKYLLEQLKERFKDKKHLDKFSSYHVKTAFFHVCTQNPQDSQWDRKDLGLCFDNCVTYFLQCLRTEKLENYFIPEFNLFSSNLIDKRSKEFLTKQIEYERNNEFPVFDEF** |
| **7** | **45bp Immunostimulatory DNA (ISD) Forward**  **5’-TACAGATCTACTAGTGATCTATGACTGATCTGTACATGATCTACA-3’** |
|  | **45bp Immunostimulatory DNA (ISD) Reverse**  **5’-TGTAGATCATGTACAGATCAGTCATAGATCACTAGTAGATCTGTA-3’** |
