## Supplementary material for "Small Molecule Properties Define Partitioning into Biomolecular Condensates": Code

# Set appropriate working directory
setwd('~/Documents/Data/221109_sabari_correlation_analysis/')

# Load data & set which data we are measuring correlations for
# We modify this script with the appropriate x and y datasets and
# change out_file_name correspondingly, then use R CMD BATCH in an sbatch
# script file, which is submitted to the SLURM workload manager on our HPC
# cluster.
pc_data <- read.csv(file='Sabari_Condensates_Drug_Correlation.csv', header=T)
x <- pc_data$Mean_cGAS
y <- pc_data$Mean_Dhh1
# Set output file name
out_file_name <- 'local_test_correlation.csv'
# Set lower limit for mean
# -1.9 works when SUMOSIM is on x-axis
# -0.9 works for SH3PRM and cGAS on x-axis
mean_lower <- -0.9
# Set the number of sampling windows and resampling iterations
# Note that 100 windows, 100 iterations, tolerance 100 requires ~6 hours of compute
# time on a 48-core HPC node
n_windows <- 100
n_iterations <- 100
stopping_tolerance <- 100

# Set environment variables for # of cores.
# We run this on a 48-core HPC node so make sure the set the NCORES environment
# variable in the shell or in the sbatch script (or equivalent for the
# workload manager on your system).
ncpu = as.numeric(Sys.getenv('NCORES'))
Sys.setenv('MC_CORES'=ncpu)
# Load parallel package
library(parallel)

# Function definitions
sample_norm_bias <- function(x, mean, sd, tol) {
  # Get number of data points
  n <- nrow(x)
  # Add an index to x for bookkeeping purposes
  x$Index <- 1:n
  # Sample with normal bias
  # Create output data.frame
  out <- data.frame(Index=NA,
                    x=NA,
                    y=NA)
  # Create vector for keeping track of fraction sampled. The stopping condition
  # for the while-loop below is when the fraction sampled is constant for a
  # given number of iterations.
  frac_sampled <- c()
  # Sample count
  i <- 1
  # Set the initial condition for the while-loop
  loop <- TRUE
  while(loop == TRUE) {
    # Get a random number with given sd, but with zero mean
    random_num <- rnorm(1, 0, sd)
    # Choose a random row from the PC data.frame x
    pc_sample <- x[sample(1:n, 1), ]
    # Sampled x PC value is mean-subtracted and compared with the absolute
    # value of the random number. The sampled value is accepted if the mean-
    # subtracted PC value is less than the absolute value of the random number.
    # This samples from x with normally distributed bias
    # We only keep the unique samples.
    if(abs(pc_sample$x - mean) <= abs(random_num) & !pc_sample$Index %in% out$Index) {
      out[i, ] <- c(pc_sample$Index, pc_sample$x, pc_sample$y)
      i <- i+1
    }
    # Calculate the fraction sampled
    frac_sampled <- c(frac_sampled, length(unique(out$Index)) / n)
    # Check the stopping condition - stop looping if the last tol iterations
    # have a constant fraction sampled
    n_samples <- length(frac_sampled)
    # Only evaluate the stopping condition if number of samples is at least tol
    if(n_samples >= tol) {
      # Get the last value in frac_sampled
      last_frac_sampled <- frac_sampled[n_samples]
      # Get the last tol samples inf frac_sampled
      tol_frac_sampled <- frac_sampled[(n_samples-tol+1):n_samples]
      # Evaluate stopping condition
      if(all(tol_frac_sampled == last_frac_sampled)) {
        loop <- FALSE
      }
    }
  }
  return(list(Samples=out, FracSampled=last_frac_sampled))
}

get_corr_in_windows_parfun <- function(mean, sd, x, y, niter, tol) {
  # Intended to be run in parallel using parallel::mcmapply
  # mean is the mean of the biasing normal distribution
  # sd is the standard devation of the biasing normal distribution
  # x is the data on the x-axis of the correlation calculation
  # y is the data on the y-axis of the correlation calculation
  # niter is the number of resampling iterations to be run

  # Make x and y into a data.frame
  d <- data.frame(x=x, y=y)
  # Do biased sampling and correlation coefficient calculation
  corrs <- rep(NA_real_, niter)
  fracs_sampled <- rep(NA_real_, niter)
  for(j in 1:niter) {
    samples <- sample_norm_bias(d, mean, sd, tol)
    corrs[j] <- cor(x=samples$Samples$x, y=samples$Samples$y)
    fracs_sampled[j] <- samples$FracSampled
  }
  # Get quartiles of the correlation coefficients calculated from resampled data
  cor_stats <- quantile(corrs, c(0, 0.25, 0.5, 0.75, 1))
  # Append the fraction sampled to cor_stats
  cor_stats <- c(cor_stats, mean(fracs_sampled))
  return(cor_stats)
}

# Run within-window correlation analysis in parallel
# Create the data.frame with desired means and standard deviations
normal_pars <- expand.grid(Mean=seq(mean_lower, 3.9, l=n_windows),
                           SD=seq(0.1, 5.5, l=n_windows))
# Calculate resampled correlation coefficients in parallel
cor_stats_par <- mcmapply(get_corr_in_windows_parfun,
                                         mean=normal_pars$Mean,
                                         sd=normal_pars$SD,
                                         MoreArgs = list(x=x,
                                                         y=y,
                                                         niter=n_iterations,
                                                         tol=stopping_tolerance))
# get_corr_in_windows_parfun() returns the quartiles of the bootstrapped
# within-window correlation coefficient. Here we just take the median,
# but to get the full set of quartiles you could run
# normal_pars$CorMin <- cor_stats_par[1, ]
# normal_pars$CorQ1 <- cor_stats_par[2, ]
# normal_pars$CorQ2 <- cor_stats_par[3, ]
# normal_pars$CorQ3 <- cor_stats_par[4, ]
# normal_pars$CorMax <- cor_stats_par[5, ]
normal_pars$Cor <- cor_stats_par[3, ]
normal_pars$FracSampled <- cor_stats_par[6, ]
# Save the data
write.csv(normal_pars, file=out_file_name,
          quote=FALSE, row.names = F)
